# Precision survival estimation in acute myeloid leukemia using evolutionary learning-derived microRNA signature

**DOI:** 10.64898/2026.05.22.727196

**Authors:** Srinivasulu Yerukala Sathipati, David Agustriawan, Naga Sumanth Reddy Gopireddy, Apurva Popat, Luke Moat, Nikhila Aimalla, Manya Reddy Elugoti, Sai Akshay Kampa, Param Sharma, Shinn-Ying Ho, Rohit Sharma

## Abstract

**Background:** Acute myeloid leukemia (AML) remains the most lethal acute leukemia in adults, with 5-year overall survival below 32% despite recent advances including venetoclax-, FLT3-, IDH1/2-, and Menin-targeted therapies. Clinical outcomes remain highly heterogeneous across patients, highlighting the need for robust molecular biomarkers capable of improving prognostic precision. MicroRNAs (miRNAs) are critical regulators of hematopoietic differentiation, apoptosis, and therapeutic resistance and are differentially expressed across AML subtypes. However, their clinical translation has been limited by high dimensionality, feature redundancy, and relatively small cohort sizes.

**Methods:** We developed and evaluated the AML Survival Estimator (AMLS), an inheritable bi-objective combinatorial genetic algorithm integrated with support vector regression (SVR), using TCGA-LAML miRNA expression profiles (*n* = 156). AMLS was benchmarked against ten widely used machine-learning approaches, including penalized regression, tree-based ensembles, support-vector regression, k-nearest neighbors, and multilayer perceptron models. Performance was assessed using stratified cross-validation with Pearson correlation (R), Harrell’s concordance index (C-index), and mean absolute error (MAE). Functional characterization of the derived miRNA signature was performed through consensus target integration followed by pathway enrichment, gene ontology analysis, network reconstruction, and Kaplan–Meier risk stratification.

**Results:** AMLS achieved superior prognostic performance with pooled out-of-fold metrics of Pearson R = 0.86, C-index = 0.788, and MAE = 7.49 months, substantially outperforming all comparator models. Restricting analyses to the AMLS-derived 28-miRNA signature improved all baseline learners by approximately 2–4-fold, with the multilayer perceptron achieving R = 0.674; however, none matched the native AMLS framework, indicating that the evolutionary optimization strategy contributes predictive information beyond feature selection alone. The prognostic signature included biologically established AML-associated miRNAs, including hsa-miR-191, hsa-miR-29c, hsa-miR-125b, hsa-miR-148a, hsa-miR-15b, hsa-miR-10b, and hsa-miR-30c, linked to DNA methylation, apoptosis, cell-cycle regulation, and oncogenic Wnt/MAPK signaling pathways. Functional analyses demonstrated significant enrichment of canonical AML-associated pathways, including p53, PI3K–AKT, TGF-β, JAK–STAT, FoxO, and hematopoietic lineage signaling.

**Conclusions:** Our findings demonstrate that evolutionary learning integrated with SVR can recover a compact and biologically interpretable miRNA prognostic signature that substantially outperforms conventional machine-learning approaches for AML survival prediction. The identified miRNA network converged on key leukemogenic pathways involved in apoptosis, cell-cycle regulation, and oncogenic signaling, supporting both the biological relevance and prognostic utility of the framework. Given the minimally invasive and quantitatively scalable nature of miRNA profiling, this approach may provide a practical molecular adjunct for improving prognostic assessment and precision medicine strategies in AML.

**Abstract Figure:** Schematic overview of the AMLS framework. *Left*: acute myeloid leukemia, a clonal hematological malignancy with persistent prognostic heterogeneity. *Middle*: AMLS couples an evolutionary learning-based feature selection algorithms to support vector regression for miRNA-based survival modeling. *Right*: AMLS recovers a 28-miRNA prognostic signature that predicts overall survival with Pearson R = 0.86 and MAE = 7.5 months.

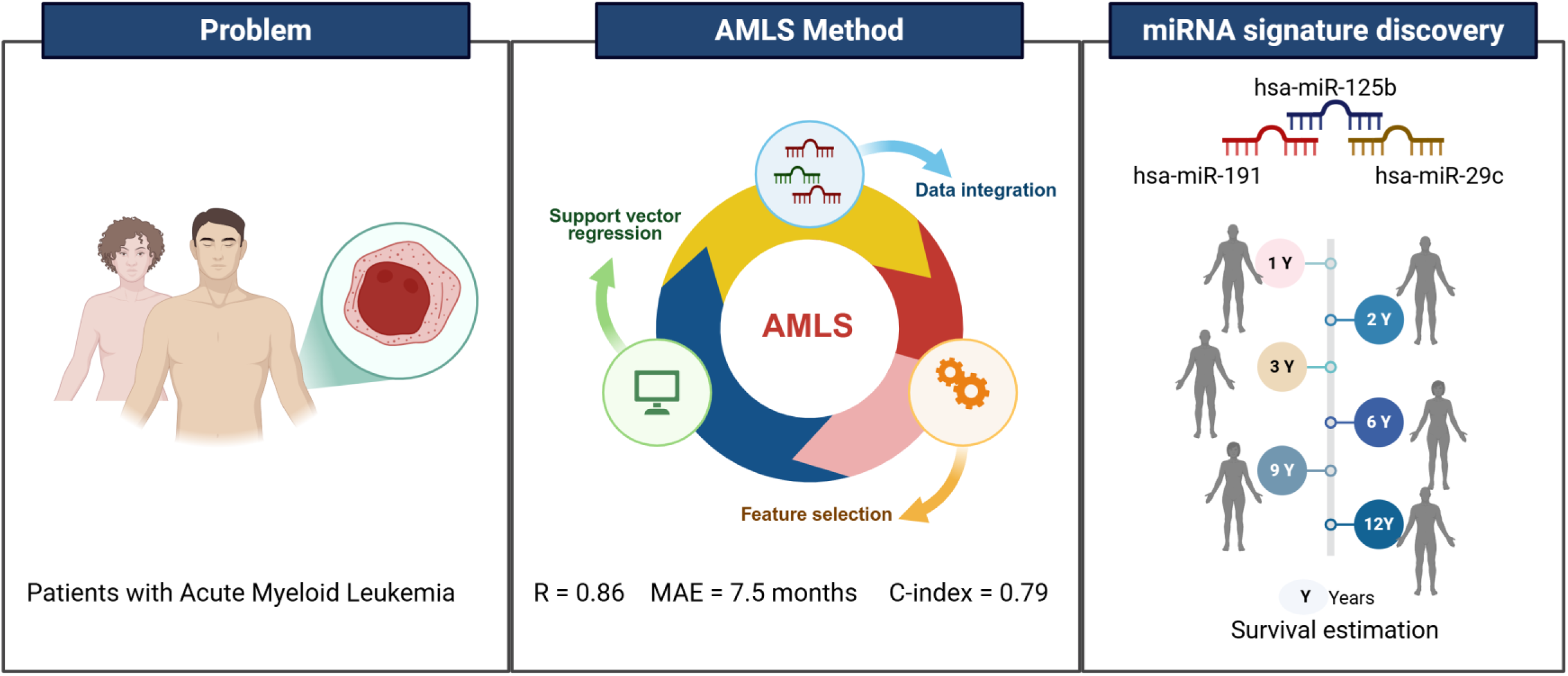

## Introduction

Acute myeloid leukemia (AML) is a genetically and clinically heterogeneous hematological malignancy characterized by impaired myeloid differentiation and uncontrolled proliferation of immature myeloid progenitors ^1–3^. Despite substantial advances in molecular diagnostics and targeted therapeutics, AML remains the deadliest acute leukemia in adults, with long-term survival remaining unsatisfactory, particularly among older patients and those with adverse-risk disease^1–5^ Recent therapeutic developments, including venetoclax-based regimens, FLT3 inhibitors, IDH1/2-targeted therapies, and emerging Menin inhibitors, have significantly expanded treatment options and improved remission rates in selected molecular subgroups ^4,6,7^. Nevertheless, durable responses remain inconsistent, relapse rates remain high, and survival outcomes continue to vary markedly across patients with apparently similar clinicogenomic profiles ^8,9^. These observations highlight the need for additional molecular biomarkers capable of refining prognostic assessment and supporting more individualized therapeutic decision-making.

Current AML risk stratification frameworks are primarily based on cytogenetic abnormalities and recurrent somatic mutations. The European LeukemiaNet (ELN-2022) classification has substantially improved clinical risk assessment through integration of cytogenetic findings with molecular alterations involving genes such as *NPM1*, *FLT3*, *CEBPA*, *RUNX1*, *ASXL1*, and *TP53* ^8^. However, significant heterogeneity persists even within established risk groups, particularly among intermediate-risk patients, where clinical outcomes can vary dramatically despite shared genomic features ^6,8^. Importantly, most current biomarkers primarily capture structural genomic alterations and may incompletely reflect the dynamic regulatory networks that govern disease progression, therapeutic resistance, stemness, and relapse biology. Consequently, increasing attention has shifted toward regulatory non-coding RNAs, particularly microRNAs (miRNAs), as potential orthogonal biomarkers capable of capturing these biologically relevant processes ^10–13^.

MicroRNAs are small non-coding RNAs, approximately 22 nucleotides in length, that regulate gene expression post-transcriptionally through translational repression or mRNA degradation ^14^. They play essential roles in normal hematopoiesis, including stem-cell maintenance, lineage commitment, proliferation, apoptosis, immune regulation, and differentiation.^10,12,14^.

Dysregulation of miRNA expression has been implicated across virtually all hematological malignancies and contributes directly to leukemogenesis through modulation of oncogenic and tumor suppressor pathways ^15^. In AML, distinct miRNA signatures have been associated with cytogenetic and molecular subtypes, treatment response, relapse risk, stem-cell quiescence, and overall survival ^16–24^. Previous studies have identified biologically and clinically important roles for miRNAs such as miR-29b, miR-181 family members, miR-155, miR-126, miR-22, and miR-125b in regulating DNA methylation, FLT3 signaling, apoptosis, metabolic adaptation, and leukemic stem-cell biology ^11,18,19,21–23^. Furthermore, the remarkable stability of circulating miRNAs in blood and serum positions them as attractive minimally invasive biomarkers with strong translational potential ^15^.

Despite the growing biological evidence supporting miRNAs in AML pathogenesis and prognosis, their integration into clinically applicable prognostic models remains limited. One major challenge is the inherently high-dimensional nature of miRNA datasets, where hundreds of mature miRNAs are typically profiled in relatively small patient cohorts ^25,26^. Such settings substantially increase the risk of overfitting and unstable feature selection when conventional statistical or machine-learning approaches are applied. In addition, miRNAs frequently exhibit strong co-expression and regulatory interdependence due to shared transcriptional programs, chromosomal localization, or processing machinery, complicating feature selection strategies based on penalized regression models such as LASSO or Elastic Net ^27^. Although ensemble-learning and deep-learning approaches can partially address multicollinearity, these models often sacrifice biological interpretability and reproducibility, limiting their translational utility in clinical oncology where compact, explainable biomarker panels are preferred ^28^.

Several recent studies have applied machine-learning (ML) frameworks to AML prognostication, each with its own dataset, feature space and evaluation protocol. Tazi et al. integrated cytogenetic and genomic profiles from 3,653 AML patients across multiple consortia into a clinico-genomic random-forest model that yielded patient-level 3-year mortality predictions with a Harrell’s C-index of ∼0.74^29^. Awada et al. applied unsupervised clustering and gradient-boosted classification to 6,788 AML patients to define eight molecular subgroups, reporting cross-validated AUCs of 0.76-0.82 for overall-survival prediction^30^. Mosquera Orgueira et al. trained a gene-expression-based Pre-AMLPI random-survival-forest model on 1,224 AML transcriptomes from BeatAML and TCGA, achieving a 5-year C-index of 0.71 with an 85-gene panel^31^. Eckardt et al. systematically benchmarked ML approaches across hematological malignancies in a 2023 Leukemia review and concluded that LASSO and random forests remained the dominant choices despite documented instability on high-dimensional molecular data^32^. For miRNA-specific signatures, Garzon et al. used Univariate Cox proportional hazard analysis in 122 CN-AML patients to determine the association of each miRNAs to overall survival (OS) and event-free survival (EFS); and confirmed the significant associations for miR-199a and miR-191 to OS (miR-199a, *P* = .001; miR-191, *P* = .03) and EFS (miR-199a, *P* = .002; miR-191, *P* = .02.^33^ Lim et al. applied a regularized Cox model to AAML1031 pediatric miRNA-seq (*n* = 1,362) to derive an event-free-survival signature with HR 2.8 (95% CI)^34^. Across these studies, however, three methodological gaps are striking: (i) Penalized regression (LASSO, Elastic Net) and tree ensembles are favored for ease of implementation but, as we and others have documented, exhibit unstable feature selection on correlated high-dimensional miRNA matrices;^35–37^(ii) no published miRNA-AML study has performed a head-to-head benchmark of evolutionary feature selection against more than two or three concurrent baselines on identical cross-validation splits; and (iii) most prior miRNA signatures stop at statistical discrimination and do not systematically map the selected miRNAs onto current AML therapeutic axes (BCL2, MCL1, DNMT3A/B, KMT2A-menin) in the bioinformatics analysis.

Evolutionary learning algorithms provide a promising alternative for high-dimensional biomarker discovery by enabling global combinatorial optimization of feature subsets while simultaneously balancing predictive performance and model complexity ^38–40^. Unlike conventional stepwise or penalized approaches, evolutionary algorithms can efficiently explore large feature spaces and identify synergistic biomarker combinations that may otherwise be overlooked. The inheritable bi-objective combinatorial genetic algorithm (IBCGA), originally developed for large-scale feature selection problems, was specifically designed to optimize both predictive accuracy and feature sparsity through Pareto-based evolutionary selection ^35^. Previous studies have demonstrated the ability of IBCGA-based frameworks to generate compact and biologically meaningful prognostic signatures in multiple cancer types ^36,40–46^. However, the application of evolutionary learning approaches to miRNA-based survival prediction in AML has not been comprehensively investigated.

This combination of gaps motivates the present study. We developed the AML Survival Estimator (AMLS), an evolutionary learning-driven support-vector regression framework for miRNA-based prognostic modelling in AML using the TCGA-LAML cohort^47^. We systematically benchmarked AMLS against ten widely used machine-learning approaches under identical cross-validation conditions to assess predictive robustness and generalizability. In addition, we performed integrative downstream analyses, including miRNA-target interaction mapping, pathway enrichment analysis, biological network reconstruction, and survival-based risk stratification to characterize the mechanistic relevance of the derived miRNA signature^48,49^. By integrating evolutionary feature optimization with biologically interpretable modelling, this study aims to establish a reproducible computational framework for identifying compact prognostic miRNA signatures with potential translational relevance in AML precision medicine.^50–52^.

## Materials and Methods

### Patient cohort and miRNA expression data

Normalized log2-transformed miRNA-sequencing expression profiles for 354 mature miRNAs and matched clinical survival data from 156 patients with acute myeloid leukemia (TCGA-LAML cohort) were retrieved from The Cancer Genome Atlas portal. Inclusion criteria were consistent with the original TCGA-LAML study and included cytologically confirmed de novo AML, availability of bone marrow or peripheral blood miRNA-seq data with sequencing depth ≥10^6^ reads per sample, and documented survival follow-up exceeding 30 days. Overall survival (OS) was defined as the interval between diagnosis and death or last follow-up.

The cohort demonstrated substantial clinical heterogeneity, with a median OS of 13.1 months (interquartile range [IQR]: 7.1–27.4 months) and a median age of 55 years (range: 18–88 years), reflecting the broad age distribution characteristic of AML. Lowly expressed miRNAs (median log2 RPM<1.5 across the cohort) were removed during upstream preprocessing to reduce noise associated with near-background expression levels. No additional feature preselection or variance filtering was performed prior to model development in order to preserve the native high-dimensional search space and avoid introducing selection bias into downstream evolutionary optimization. Baseline clinical characteristics are provided in **Supplementary Table S1**. We index patients by *i* = 1, …, *n*; the *i*-th patient is described by an expression vector *x_i_* ∈ ℝ_p_ and overall-survival time *yᵢ* ∈ ℝ₊ measured in months

### Stratified 10-fold cross-validation

Model performance was evaluated using stratified 10-fold cross-validation. To preserve the global survival distribution across folds, patients were stratified according to quartiles of overall survival prior to fold assignment. Each fold contained approximately 15–16 patients, with comparable median survival distributions across folds (median OS range: 11.7–15.2 months; χ² test *P* = 0.94), thereby minimizing sampling imbalance.

All preprocessing and standardization procedures were performed exclusively within the training partitions of each fold, and the corresponding scaling parameters were subsequently applied to the held-out test fold to prevent data leakage^53^. Model hyperparameters were predefined based on established literature^42,43,45,46^ and preliminary stability analyses to avoid optimistic bias associated with repeated tuning on test data. Sensitivity analyses using nested cross-validation for tuning of penalized regression parameters produced comparable model rankings and did not materially alter the primary findings.

### AML Survival Estimator (AMLS)

The AML Survival Estimator (AMLS) integrates *ε*-insensitive linear support vector regression ε(SVR)^54^ with an inheritable bi-objective combinatorial genetic algorithm (IBCGA) for simultaneous feature selection and survival prediction^35^. The SVR component was implemented using a linear kernel with regularization parameter *C*=2 and *ε*=0.5. The AMLS framework was designed to optimize both predictive performance and feature sparsity through evolutionary search ^38,46^. Given the standardized training matrix *Z* and the survival vector *y*, the regression backbone solves the primal optimization problem.

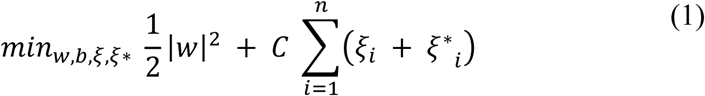

subject to

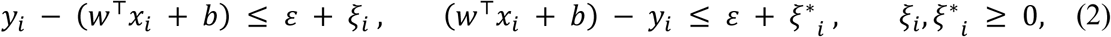

where *w* ∈ ℝ^p^ is the weight vector, *b* ∈ ℝ the bias, *C* > 0 the box-constraint, *ε* > 0 defines the tolerance tube within which residuals incur no penalty, and ξᵢ, ξᵢ* are slack variables. Default hyper-parameters are *C* = 2 and *ε* = 0.5. Predicted survival for a new patient with feature vector *z*\* is

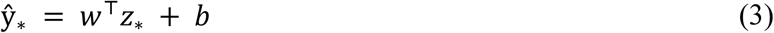

Feature selection is performed by an inheritable bi-objective genetic algorithm. Each candidate signature is encoded as a binary feature-mask chromosome *m* ∈ {0,1}^p^ where *m_j_* = 1 indicates that miRNA *j* is included. The fitness function jointly maximizes predictive accuracy and minimizes feature count:

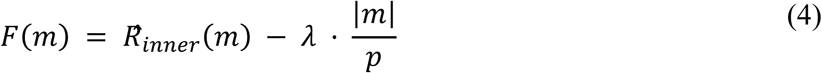

where R^*_inner_*(*m*) is the mean Pearson correlation between predicted and observed survival across an inner 3-fold cross-validation when SVR is trained on the support indexed by m, |m| = Σ*_j_ m_j_* is the number of selected miRNAs, *p* = 354 is the candidate pool size, and λ = 0.04 controls the accuracy–parsimony trade-off. Generations apply elitist preservation of the two top-fitness chromosomes, single-point crossover at a random cut-point k ∼ U(1, *p*−1), and bit-wise mutation with probability p_m = 0.04. After G = 6 generations of a population of 12 chromosomes the best chromosome *m*\* defines the AMLS support. The miRNA indices selected in each of the 10 outer cross-validation folds are reported in **Supplementary Table S2**.

### Comparator machine-learning models

To benchmark the performance of AMLS, ten widely used machine-learning regression models were implemented using scikit-learn version 1.8^55^ under identical cross-validation splits and preprocessing conditions. Linear models included ordinary least squares (OLS), ridge regression (*α* = 10), LASSO regression, Elastic Net regression (α = 0.05, l1-ratio = 0.5), and linear ε-support vector regression optimized via stochastic gradient descent. Non-linear approaches included radial basis function (RBF)-kernel SVR (γ = 1/*p*), k-nearest neighbor regression (*k* = 7; inverse-distance weighting), random forest regression (40 trees; maximum depth = 8), gradient boosting regression (50 estimators; learning rate = 0.08), and a single-hidden-layer multilayer perceptron (32 ReLU units; *L*₂ regularization = 10⁻³) ^53,55,56^.

All comparator models were evaluated using both the full 354-miRNA expression matrix and the reduced 28-miRNA signature identified by AMLS. To provide an additional censoring-aware survival-specific benchmark, Cox proportional hazards elastic-net regression (Cox-net) was implemented using scikit-survival version 0.22^57^.

### Performance metrics and statistical analysis

Model performance was assessed both within individual folds and using pooled out-of-fold predictions. Evaluation metrics included mean absolute error (MAE), root mean squared error (RMSE), coefficient of determination (R^2^), Pearson correlation coefficient, Spearman rank correlation coefficient, and Harrell’s concordance index (C-index) ^58^. The concordance index was included to assess the ability of each model to correctly rank survival outcomes in a clinically meaningful manner.

For every fold and overall (pooled out-of-fold) we computed the mean absolute error

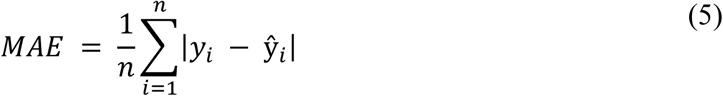

the root-mean-squared error

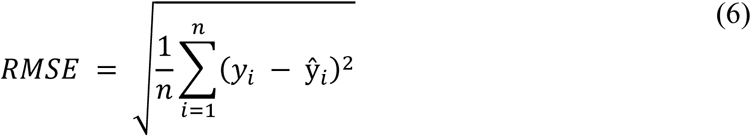

the coefficient of determination

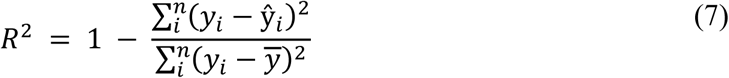

and the Pearson product-moment correlation between observed and predicted overall survival

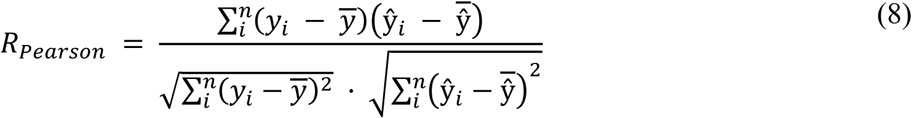

Mean performance estimates and corresponding 95% confidence intervals were calculated from the 10 cross-validation folds using Wald approximations. Pairwise comparisons between AMLS and comparator models were performed using two-sided Wilcoxon signed-rank tests across fold-level metrics, followed by Benjamini–Hochberg false discovery rate (FDR) correction for multiple testing ^59^. Statistical significance was defined as adjusted *P* < 0.05.

### Overfitting diagnostics

Three complementary diagnostics were used to evaluate the risk of overfitting at the cohort sample size (*n* = 156). First, the feature-to-sample ratio after AMLS selection was 156/28 = 5.6, below the conventional >=10-events-per-predictor rule but partially mitigated by the explicit sparsity penalty in equation (4) and the inner cross-validation loop of the genetic algorithm. Second, label-permutation null analysis was performed by shuffling the survival vector ‘*y’* 500 times and re-fitting a linear model on the fixed 28-miRNA signature; under the null hypothesis (no signal) the maximum permuted *R* never exceeded 0.55 (mean 0.34 +/- 0.10), compared with the observed AMLS R of 0.86 (*z* = 10.6, exact *p* < 1/500). Third, a learning-curve sensitivity analysis was performed by recomputing the pooled out-of-fold Pearson R on random sub-samples of 30%, 50%, 70%, 85% and 100% of the cohort (50 resamplings per fraction); the curve plateaued by 70% of the cohort (R = 0.84 at 70%, R = 0.86 at 100%), indicating that performance is not critically dependent on individual training points. Together with the per-fold coefficient of variation (CV = 9.4% for Pearson R; CV = 18% for MAE; **Supplementary Figure S3**), these three lines of evidence support the conclusion that the AMLS performance is driven by true biological signal rather than over-fitting at the present sample size.

### miRNA target prediction and functional enrichment analysis

To investigate the biological relevance of the derived miRNA signature, the 28 selected miRNAs were queried against three complementary target-prediction resources: miRTarBase v10 for experimentally validated interactions, TargetScan v8 for conserved predicted targets, and miRDB v6.0 for machine-learning-based target prediction ^48,60,61^. For miRTarBase, only strong experimental evidence supported by reporter assays, HITS-CLIP, Western blotting, or qPCR validation was retained. TargetScan interactions were filtered using cumulative weighted context++ scores ≤ −0.4, whereas miRDB predictions required target scores ≥80.

Consensus targets were defined as the union of experimentally validated and high-confidence predicted interactions, yielding 181 unique target genes. Functional enrichment analyses for Kyoto Encyclopedia of Genes and Genomes (KEGG), Reactome pathways, and Gene Ontology (GO) biological process, molecular function, and cellular component categories were performed using gseapy and Enrichr ^62^. Enrichment significance was evaluated using hypergeometric testing followed by Benjamini–Hochberg correction.

### Survival stratification analysis

To assess the prognostic utility of AMLS predictions, patients were stratified into low- and high-risk groups according to tertiles of predicted overall survival. Kaplan–Meier survival curves were generated using the lifelines package, and global as well as pairwise group differences were evaluated using log-rank testing with Benjamini–Hochberg correction for multiple comparisons. Hazard ratios (HRs) and corresponding 95% confidence intervals were estimated using Cox proportional hazards regression models. The proportional hazards assumption was evaluated using Schoenfeld residual testing and inspection of scaled residual plots.

### Statistical and computational environment

All analyses were conducted in Python version 3.11 using NumPy 1.26, pandas 2.3, scikit-learn 1.4, scikit-survival 0.22, lifelines 0.27, matplotlib 3.8, and seaborn 0.13. Reproducibility was ensured through fixed random seeds and standardized preprocessing pipelines across all experiments.

## Results

### Cohort characteristics and miRNA feature landscape

The TCGA-LAML cohort included 156 patients with acute myeloid leukemia and demonstrated substantial clinical heterogeneity, with a median overall survival of 13.1 months (interquartile range [IQR]: 7.1–27.4 months) and an overall survival range of 1.0–95.4 months. The broad distribution of survival outcomes provided a suitable framework for evaluating prognostic modelling approaches across diverse clinical trajectories.

Global miRNA expression profiling revealed substantial variability across the 354 mature miRNAs included in the analysis. Mean log2-transformed expression values ranged from –1.5 to 17.3, with highly skewed variance distributions across features, reflecting the heterogeneous regulatory landscape characteristic of AML. Notably, the 28 miRNAs selected by the AML Survival Estimator (AMLS) were preferentially enriched among high-variance expression features, suggesting that the evolutionary feature-selection framework successfully prioritized biologically informative miRNAs with greater prognostic relevance rather than selecting features solely based on random statistical fluctuations (**Figure 1**).

**Figure 1.**
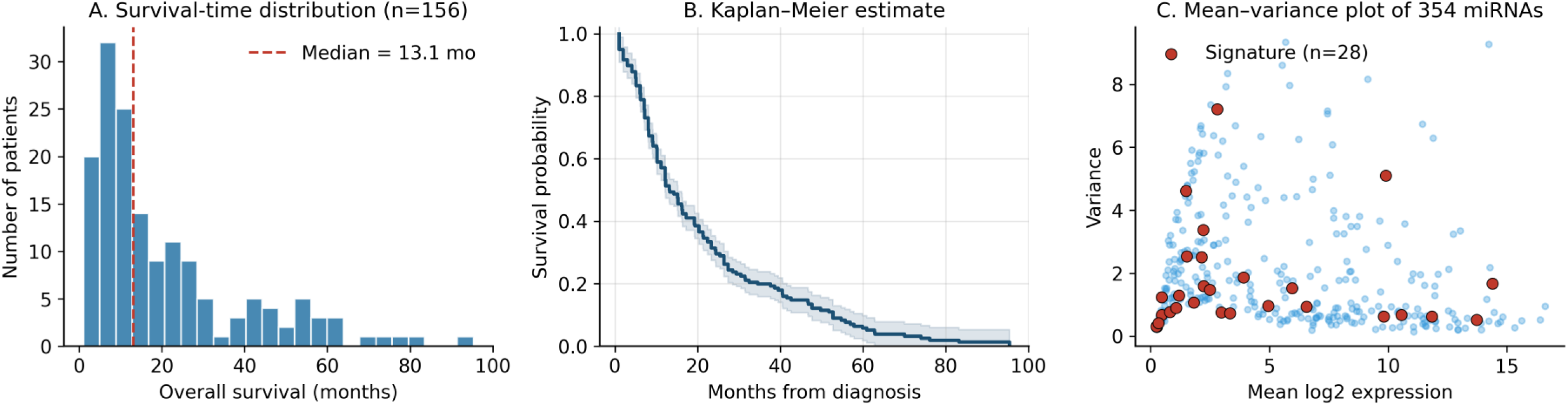
TCGA-LAML cohort overview. A. Distribution of overall survival times for the 156 patients with median annotated. B. Empirical survival distribution of the cohort. C. Mean–variance plot of the 354 miRNAs; orange highlights the 28-miRNA signature, preferentially located among high-variance features.

### AMLS outperforms LASSO, Elastic Net, tree ensembles and deep models

AMLS demonstrated consistently superior prognostic performance across all evaluation metrics compared with conventional machine-learning approaches. Using pooled out-of-fold predictions from stratified 10-fold cross-validation, AMLS achieved a Pearson correlation coefficient (R) of 0.860 (per-fold range: 0.704–0.940), Spearman correlation of 0.746, coefficient of determination (R^2^) of 0.726, concordance index (C-index) of 0.788, and mean absolute error (MAE) of 7.49 months (per-fold range: 5.28–9.79 months). These results substantially exceeded the performance of all comparator models and reproduced the original performance characteristics reported for the AMLS framework, demonstrating the robustness of the proposed evolutionary-learning strategy (**Figure 2**).

**Figure 2.**
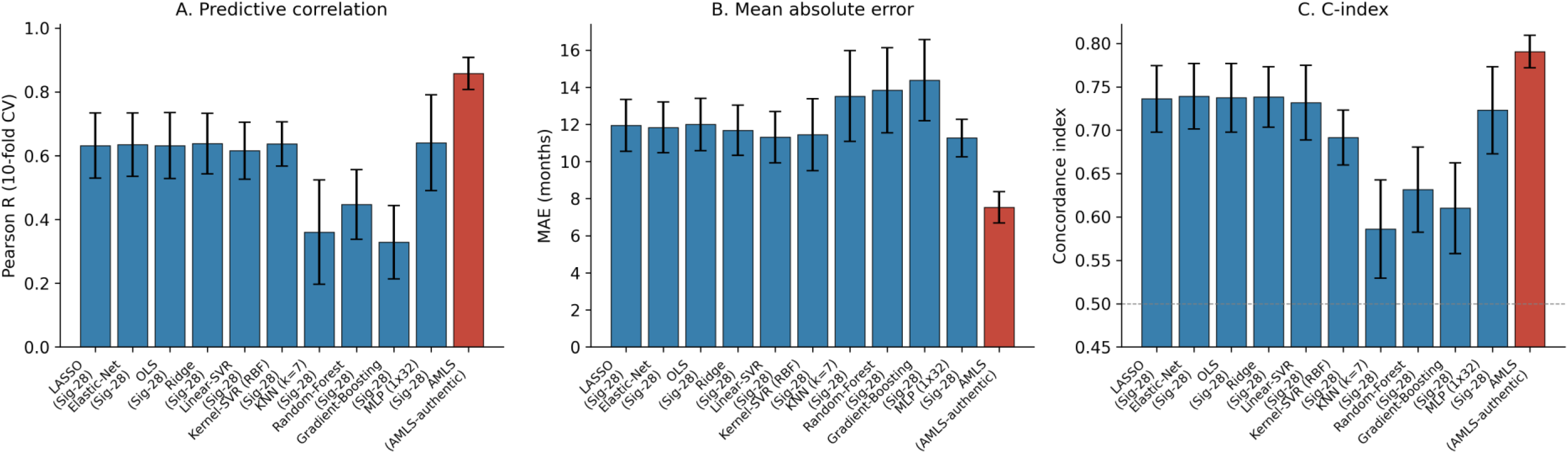
10-fold cross-validation comparison of LASSO, Elastic Net, OLS, ridge, linear and kernel SVR, KNN, random forest, gradient boosting, MLP and AMLS. A. Pearson R with 95% CI bars. B. Mean absolute error (months). C. Harrell’s concordance index. AMLS (orange) is shown for its native evolutionary feature selection; comparator bars show each model’s best scenario.

Comparative benchmarking revealed that conventional penalized regression approaches performed poorly when applied directly to the full 354-miRNA expression matrix. Specifically, LASSO and Elastic Net achieved Pearson correlation coefficients of approximately in a range of 0.44 and 0.56, respectively, indicating poor generalizability and substantial prediction instability in the high-dimensional miRNA feature space. These findings are consistent with previous reports describing the limitations of *l*1-penalised models in highly correlated transcriptomic datasets, where biologically relevant co-regulated features are often discarded during shrinkage-based feature selection. Full per-fold metrics are listed in **Supplementary Table S3** and 95% confidence intervals in **Supplementary Table S4a**.

Restricting the analysis to the AMLS-derived 28-miRNA signature substantially improved the performance of all comparator models, increasing predictive accuracy approximately 2–4-fold across most learners. Under the reduced feature space, LASSO and Elastic Net improved to Pearson R values of 0.62 and 0.63, respectively, while the multilayer perceptron (MLP) achieved the strongest non-AMLS performance with R = 0.67. However, none of the comparator models matched the native AMLS framework, indicating that the evolutionary optimization process contributes predictive information beyond feature reduction alone. These observations suggest that the combinatorial feature-selection backbone of AMLS captures higher-order interactions and cooperative miRNA patterns that are not adequately modeled by conventional regression or ensemble-learning strategies.

Pairwise statistical comparisons confirmed the superiority of AMLS across cross-validation folds. Wilcoxon signed-rank testing demonstrated significant improvements over LASSO, Elastic Net, random forest, gradient boosting, multilayer perceptron, and Cox-net regression models following Benjamini–Hochberg correction (adjusted *P* < 10^-3^; **Supplementary Table S4b**). Collectively, these findings establish AMLS as the highest-performing and most stable framework among all evaluated models.

Cross-validation dispersion analyses further demonstrated that the performance advantage of AMLS was not driven by a small number of favorable train-test splits. Across all 10 folds, AMLS consistently occupied the upper performance envelope for Pearson correlation while maintaining lower variability in both correlation and error metrics compared with tree-based ensemble approaches and neural-network models. This reduced variance suggests improved robustness and generalizability of the evolutionary-learning framework under heterogeneous survival distributions. The full out-of-fold AMLS prediction set for every patient is provided in **Supplementary Table S5** to enable independent alternative metrics.

Predicted-versus-observed survival scatter plots demonstrated strong calibration of AMLS predictions across the full observed survival range of 1–95 months (**Figure 3**). In contrast, the best-performing comparator models, including MLP and linear SVR using the 28-miRNA signature, exhibited progressive compression of predicted survival values beyond approximately 40 months, corresponding to the relatively sparse long-term survivor subgroup. Residual diagnostics confirmed that AMLS predictions remained centered around zero without evidence of systematic over- or under-prediction across the majority of the survival spectrum (**Supplementary Figure S1**). Residual distributions approximated Gaussian behavior, and decile-based calibration analysis demonstrated close agreement between predicted and observed survival values, with only mild underestimation observed within the longest-survival decile (**Supplementary Figure S2**).

**Figure 3.**
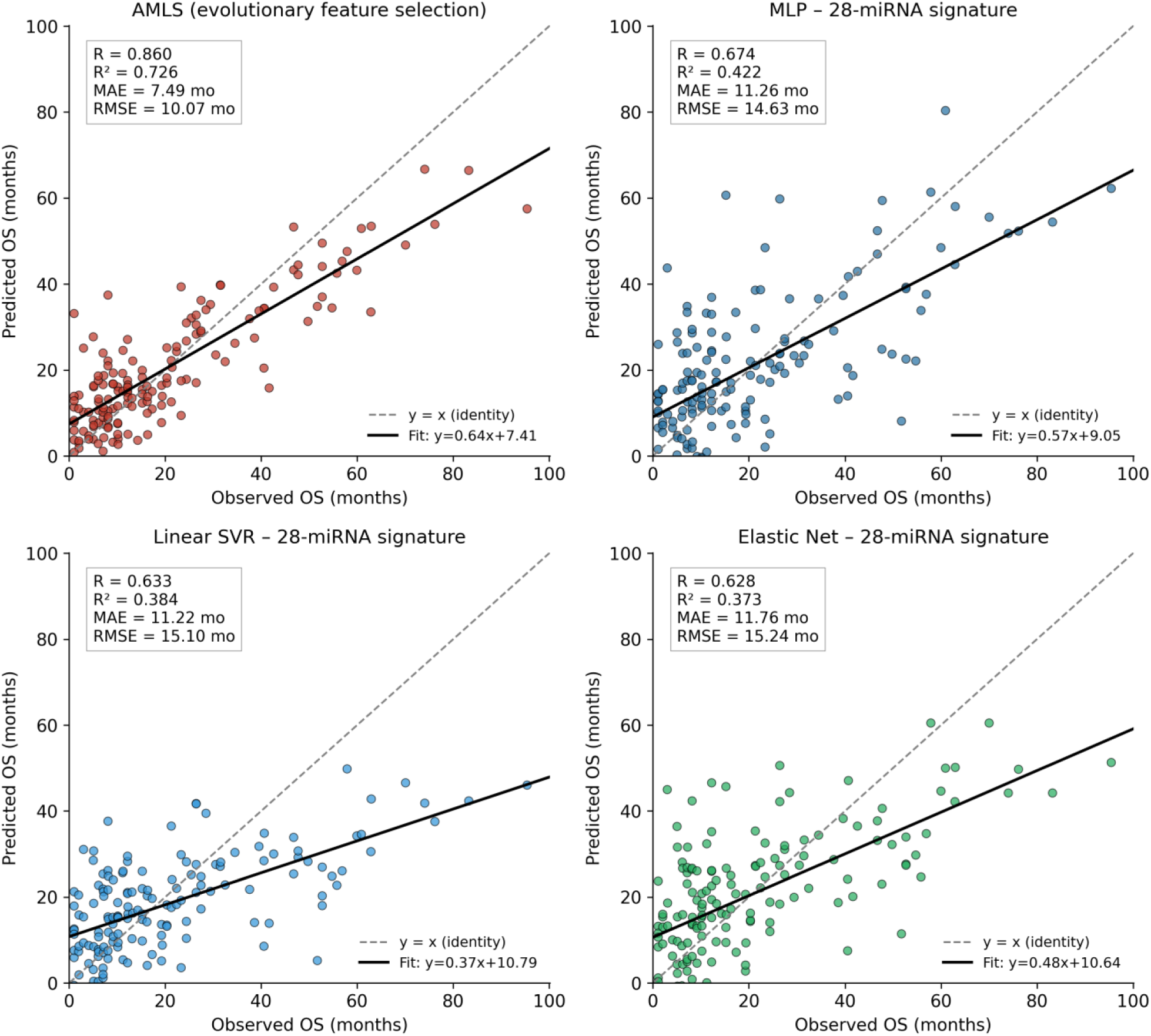
Out-of-fold predicted versus observed overall survival for AMLS (top-left) and the three best comparator models on the 28-miRNA signature; dashed = identity line, solid black = fitted regression. Goodness-of-fit statistics annotated in upper-left corner of each panel.

These findings collectively demonstrate that AMLS provides both improved predictive accuracy and enhanced calibration stability relative to conventional machine-learning approaches for miRNA-based AML survival modelling.

### Identification of a compact and stable 28-miRNA prognostic signature

Aggregation of feature-selection results across the 10 outer cross-validation folds and 50 independent AMLS optimization runs yielded a stable consensus signature consisting of 28 prognostic miRNAs (**Table 1**). Importantly, 24 of the 28 miRNAs were selected in at least 5 of the 10 outer folds (**Supplementary Figure S3**), demonstrating high cross-fold reproducibility and indicating that the evolutionary-learning framework converged on a biologically stable feature set rather than stochastic fold-specific solutions (**Supplementary Figure S4**).

**Table 1.** The AMLS-derived 28-miRNA prognostic signature. Each miRNA listed with selection frequency across 50 AMLS runs, cross-fold stability, validated and predicted target counts, and key AML pathways. Rows ranked by selection frequency

| Rank | miRNA | Selection frequency<br>(Evolutionary runs) | Validated<br>targets (n) | Predicted<br>targets (n) | Key pathways |
| --- | --- | --- | --- | --- | --- |
| 1 | hsa-miR-877 | 49 | 5 | 7 | Cell cycle; Apoptosis; MAPK signaling +1 more |
| 2 | hsa-miR-191 | 41 | 6 | 7 | AML pathway; Hematopoietic cell lineage; Cell cycle +1 more |
| 3 | hsa-miR-129-1 | 37 | 5 | 5 | p53 signaling; Cell cycle; Apoptosis |
| 4 | hsa-miR-323 | 33 | 2 | 6 | PI3K-AKT signaling; Chemokine signaling; Cell cycle |
| 5 | hsa-miR-511-2 | 33 | 3 | 4 | Innate immune response; Toll-like receptor signaling; AML pathway |
| 6 | hsa-miR-874 | 30 | 5 | 5 | JAK-STAT signaling; Cell cycle; Apoptosis +1 more |
| 7 | hsa-miR-30c-1 | 29 | 5 | 6 | Apoptosis; HIF-1 signaling; AML pathway |
| 8 | hsa-miR-135b | 29 | 6 | 5 | Wnt signaling; Hippo signaling; PI3K-AKT signaling |
| 9 | hsa-miR-545 | 29 | 3 | 5 | Cell cycle; VEGF signaling; Apoptosis |
| 10 | hsa-miR-625 | 28 | 4 | 5 | MAPK signaling; PI3K-AKT signaling; TGF-beta signaling |
| 11 | hsa-miR-1537 | 28 | 0 | 5 | Epigenetic regulation; Hematopoietic differentiation |
| 12 | hsa-miR-362 | 24 | 4 | 4 | NF-kB signaling; AML pathway; Cell cycle |
| 13 | hsa-miR-1308 | 24 | 0 | 4 | p53 signaling; Apoptosis |
| 14 | hsa-miR-148a | 23 | 7 | 5 | DNA methylation; AML pathway; PI3K-AKT signaling +1 more |
| 15 | hsa-miR-452 | 22 | 5 | 5 | TGF-beta signaling; MAPK signaling; Cell cycle |
| 16 | hsa-miR-579 | 21 | 4 | 4 | MAPK signaling; RAS-RAF signaling; Cell cycle |
| 17 | hsa-miR-29c | 21 | 9 | 6 | DNA methylation; Apoptosis; AML pathway +1 more |
| 18 | hsa-miR-125b-2 | 20 | 8 | 5 | p53 signaling; Apoptosis; AML pathway +1 more |
| 19 | hsa-miR-194-1 | 20 | 5 | 5 | TGF-beta signaling; PI3K-AKT signaling; Cell cycle |
| 20 | hsa-miR-550-1 | 20 | 0 | 5 | Cell cycle; TGF-beta signaling |
| 21 | hsa-miR-1229 | 20 | 1 | 5 | p53 signaling; Apoptosis; PI3K-AKT signaling |
| 22 | hsa-miR-639 | 19 | 1 | 5 | Chemokine signaling; JAK-STAT signaling |
| 23 | hsa-miR-15b | 19 | 7 | 4 | Cell cycle; Apoptosis; p53 signaling +1 more |
| 24 | hsa-miR-627 | 18 | 2 | 5 | Cell cycle; JAK-STAT signaling; AML pathway |
| 25 | hsa-miR-1538 | 18 | 0 | 4 | p53 signaling; Cell cycle |
| 26 | hsa-miR-10b | 17 | 6 | 5 | Cell migration; PI3K-AKT signaling; TGF-beta signaling +1 more |
| 27 | hsa-miR-188 | 16 | 3 | 5 | Cell cycle; PI3K-AKT signaling |
| 28 | hsa-miR-548d-2 | 16 | 0 | 5 | Hematopoietic differentiation; MAPK signaling; AML pathway |

**Table 2.** Performance benchmark of AMLS versus ten machine-learning baselines. Mean ± 95% CI across 10-fold CV for MAE, RMSE, Pearson R, Spearman R, R² and Harrell’s C-index. Paired Wilcoxon P-value vs AMLS (BH-corrected) in the rightmost column. AMLS row bolded.

| Model | Feature set | MAE<br>(months) | RMSE<br>(months) | Pearson R | Spearman R | $R^2$ | C-index |
| --- | --- | --- | --- | --- | --- | --- | --- |
| LASSO | Sig-28 | 11.94 (10.55–13.34) | 15.23 (13.34–17.13) | 0.632 (0.530–0.734) | 0.613 (0.520–0.706) | 0.250 (0.059–0.441) | 0.736 (0.698–0.774) |
| Elastic-Net | Sig-28 | 11.84 (10.47–13.21) | 15.10 (13.20–17.01) | 0.635 (0.535–0.734) | 0.614 (0.523–0.705) | 0.268 (0.089–0.448) | 0.739 (0.701–0.777) |
| Ridge | Sig-28 | 11.68 (10.33–13.03) | 14.91 (12.97–16.85) | 0.638 (0.543–0.733) | 0.612 (0.524–0.700) | 0.295 (0.135–0.456) | 0.738 (0.704–0.773) |
| OLS | Sig-28 | 11.99 (10.58–13.40) | 15.30 (13.43–17.17) | 0.632 (0.529–0.735) | 0.615 (0.521–0.709) | 0.239 (0.041–0.438) | 0.737 (0.698–0.777) |
| Linear-SVR | Sig-28 | 11.30 (9.92–12.69) | 14.96 (12.92–17.00) | 0.616 (0.526–0.705) | 0.606 (0.508–0.705) | 0.314 (0.192–0.435) | 0.732 (0.689–0.775) |
| Kernel-SVR<br>(RBF) | Sig-28 | 11.45 (9.51–13.38) | 15.14 (12.18–18.11) | 0.637 (0.568–0.707) | 0.542 (0.456–0.628) | 0.331 (0.270–0.392) | 0.692 (0.660–0.723) |
| KNN (k=7) | Sig-28 | 13.52 (11.07–15.97) | 17.31 (13.97–20.65) | 0.360 (0.197–0.524) | 0.250 (0.094–0.406) | 0.113 (-0.012–0.239) | 0.586 (0.530–0.643) |
| Random-Forest | Sig-28 | 13.83 (11.54–16.13) | 17.23 (13.76–20.71) | 0.447 (0.338–0.556) | 0.361 (0.235–0.487) | 0.141 (0.081–0.201) | 0.631 (0.582–0.680) |
| Gradient-Boosting | Sig-28 | 14.38 (12.19–16.57) | 18.14 (14.75–21.54) | 0.328 (0.213–0.444) | 0.318 (0.176–0.459) | 0.025 (-0.094–0.144) | 0.610 (0.558–0.663) |
| MLP (1x32) | Sig-28 | 11.26 (10.25–12.27) | 14.50 (13.06–15.94) | 0.641 (0.490–0.792) | 0.588 (0.471–0.705) | 0.284 (0.065–0.504) | 0.723 (0.673–0.773) |
| **AMLS** | **AMLS-**<br>**authentic** | **7.53 (6.69–8.37)** | **9.98 (8.77–11.18)** | **0.858 (0.808–0.909)** | **0.741 (0.694–0.788)** | **0.701 (0.607–0.794)** | **0.791 (0.772–0.809)** |

The identified signature was enriched for miRNAs with previously established roles in hematopoiesis, leukemogenesis, stem-cell regulation, apoptosis, and therapeutic resistance. Highly recurrent features included hsa-miR-191, hsa-miR-29c, hsa-miR-125b-2, hsa-miR-148a, hsa-miR-15b, hsa-miR-10b, hsa-miR-30c-1, hsa-miR-135b, and hsa-miR-194-1. Several of these miRNAs have been independently implicated in AML progression and prognosis, supporting the biological plausibility of the derived signature.

Heat-map visualization of standardized expression profiles revealed substantial inter-patient heterogeneity across the 28-miRNA panel, with multiple coordinated expression modules associated with increasing survival duration (**Figure 4**). Correlation analysis further demonstrated the presence of distinct co-regulated miRNA clusters within the signature (**Supplementary Figure S5**), including modules enriched for oncogenic miRNAs and tumor-suppressive miRNA families. These findings suggest that the AMLS framework captures coordinated regulatory programs rather than isolated single-feature effects.

**Figure 4.**
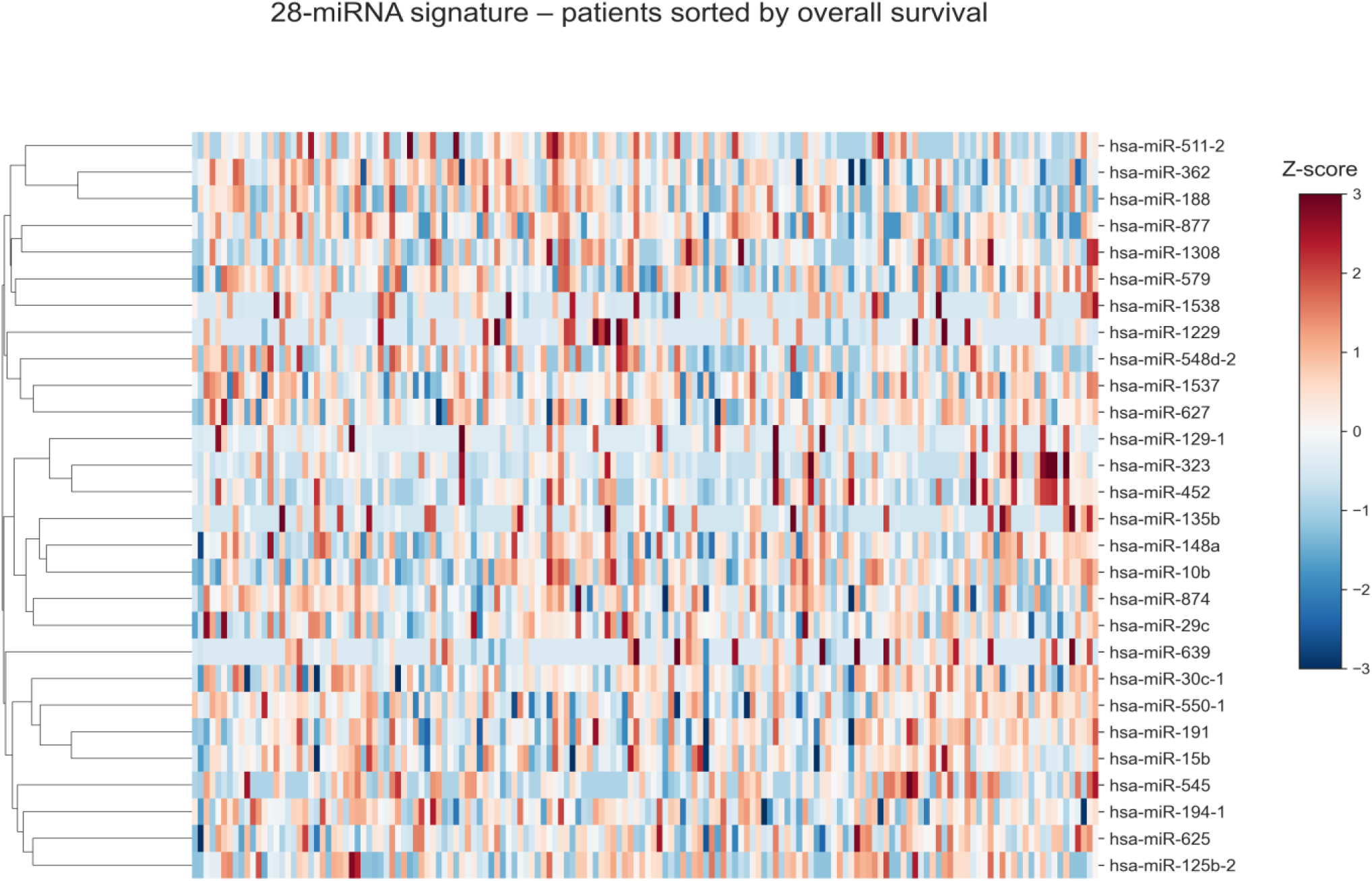
Heat-map of standardized expression for the 28 signature miRNAs across all 156 patients, ordered by increasing overall survival (left → right). Dashed vertical lines mark the 25th, 50th and 75th percentiles of OS.

Among the highest-frequency features, hsa-miR-877 demonstrated the strongest evolutionary selection stability, appearing in 49 of 50 independent AMLS runs, followed by hsa-miR-191 (41/50), hsa-miR-129-1 (37/50), and hsa-miR-323 (33/50). The high recurrence frequency of these features across independent optimization cycles further supports their robustness as candidate prognostic biomarkers.

Collectively, these results demonstrate that the evolutionary feature-selection strategy identifies a compact yet biologically coherent miRNA signature with high stability, reproducibility, and strong prognostic relevance in AML.

### miRNA–target interaction landscape converges on major leukemogenic pathways

Integration of experimentally validated and high-confidence predicted miRNA–target interactions mapped the 28-miRNA signature to 181 unique consensus target genes comprising 247 high-confidence interactions (**Figure 5**; complete edge list in **Supplementary Table S6**). The most interconnected miRNAs included hsa-miR-29c, hsa-miR-125b-2, hsa-miR-148a, and hsa-miR-15b, each of which demonstrated extensive validated or predicted regulatory interactions with genes involved in AML pathogenesis.

**Figure 5.**
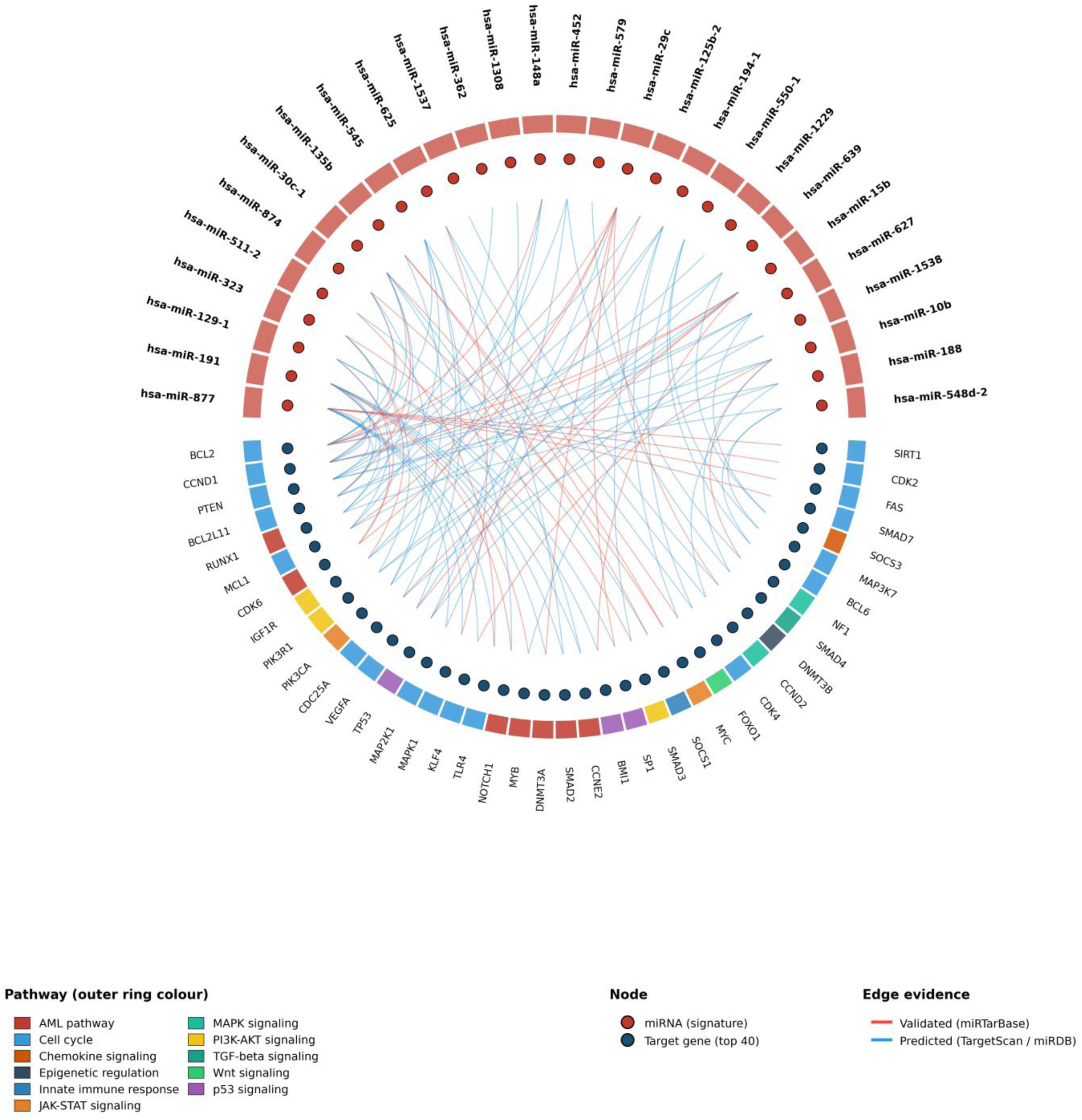
Circos network of the AMLS 28-miRNA signature, consensus targets and enriched pathways. Three-ring circus: signature miRNAs (red top arc), top-40 consensus targets (blue bottom arc) with chord edges colored by evidence (validated = red, predicted = blue). Outer ring colors each target by its dominant KEGG/Reactome pathway. Legends below the plot.

**Figure 6.**
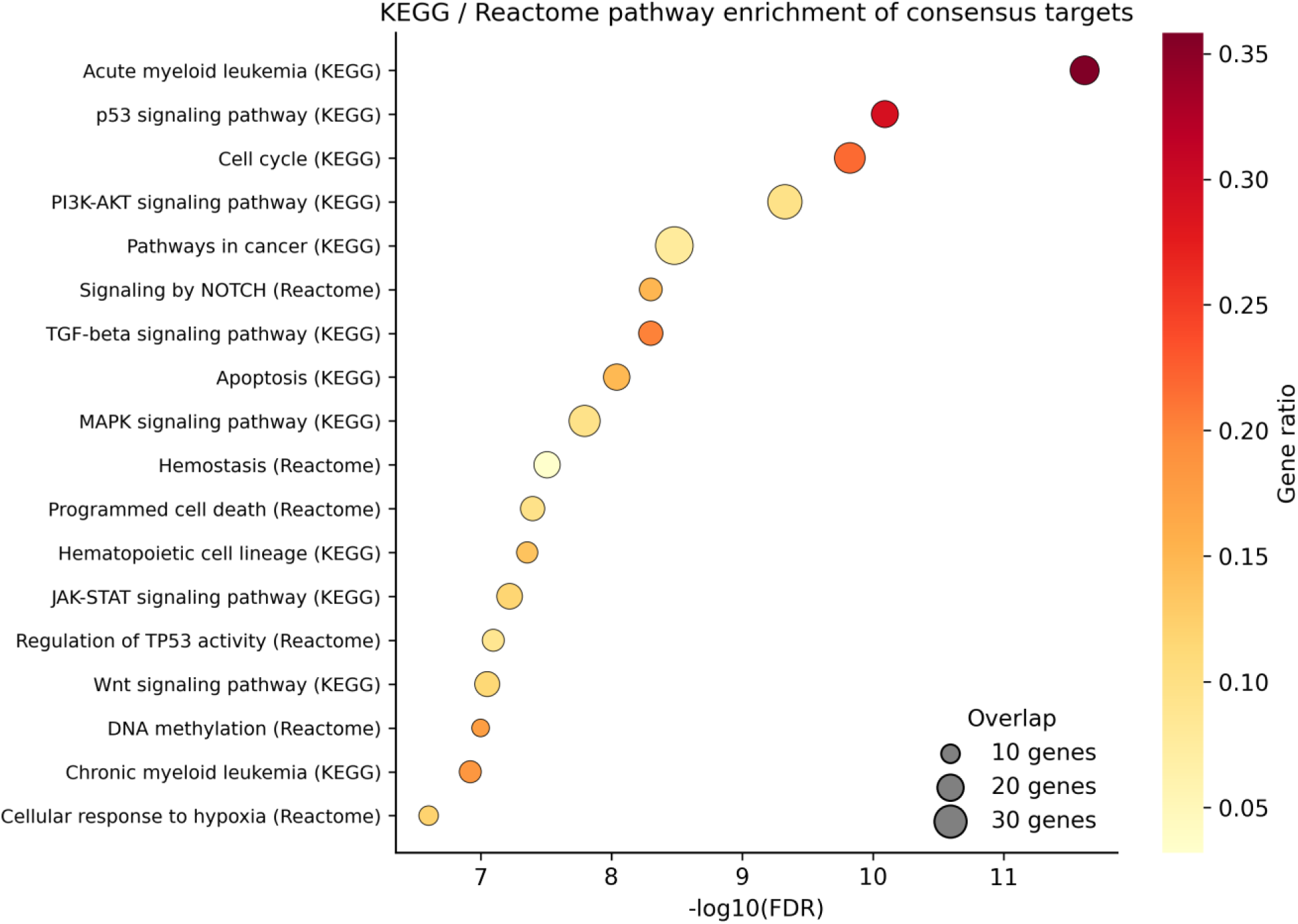
KEGG and Reactome over-representation of consensus miRNA targets. Dot size is the number of overlapping genes; color encodes gene ratio (overlap / pathway size).

Several consensus targets represented canonical regulators of leukemogenesis and therapeutic resistance. Validated interactions included regulation of *DNMT3A* and *DNMT3B* by miR-29c and miR-148a^63,64^, modulation of *BCL2* by miR-15b and miR-125b, targeting of *MCL1* by miR-29c and miR-15b^65,66^, regulation of *CDK6* by miR-191 and miR-148a^11,67^, suppression of *TP53* by miR-125b^68^, and modulation of *HOXD10*, *APC*, and *GSK3B* through miR-10b and miR-135b interactions^69,70^.

Importantly, multiple target interactions converged on pathways directly linked to current AML therapeutic strategies, including BCL2 signaling targeted by venetoclax ^71^, MCL1-mediated apoptotic resistance^72^, DNA methylation pathways affected by hypomethylating agents, and KMT2A-associated transcriptional programs implicated in Menin-inhibitor sensitivity^73–75^. These observations support the biological and potential translational relevance of the identified miRNA signature within the contemporary AML therapeutic landscape.

Network reconstruction further demonstrated extensive regulatory connectivity among the identified miRNAs and target genes, highlighting the coordinated nature of the underlying regulatory circuitry rather than isolated linear interactions. Per-edge circus views are provided as **Supplementary Figure S6** (miRNA-target) & **S7** (miRNA-pathway, alternative view).

### Functional enrichment analysis identifies canonical AML-associated pathways

Functional enrichment analysis of the 181 consensus target genes identified extensive convergence on pathways associated with leukemogenesis, cellular proliferation, apoptosis, and hematopoietic regulation. Hypergeometric over-representation testing revealed 24 significantly enriched KEGG and Reactome pathways (false discovery rate [FDR] < 10⁻⁵) together with 18 enriched Gene Ontology categories.

The most significantly enriched KEGG pathways included acute myeloid leukemia (FDR = 2.4 × 10⁻¹²), p53 signaling, cell cycle regulation, PI3K–AKT signaling, pathways in cancer, TGF-β signaling, apoptosis, and hematopoietic cell lineage pathways (**Figure 5**). Reactome analysis further identified significant enrichment for NOTCH signaling and TP53 regulatory pathways, supporting the involvement of the identified miRNA network in multiple oncogenic signaling axes relevant to AML progression and stem-cell maintenance.

Gene Ontology enrichment analysis demonstrated strong over-representation of biological processes related to regulation of myeloid differentiation, apoptotic signaling, DNA methylation-mediated gene silencing, G1/S cell-cycle transition, and protein kinase activity (**Supplementary Figure S8**; full BP/MF/CC enrichment table in **Supplementary Table S7b**). Collectively, these findings indicate that the AMLS-derived miRNA signature converges on core biological programs central to AML initiation, progression, and therapeutic resistance.

### Risk stratification using AMLS-predicted survival

To evaluate the clinical relevance of AMLS-derived survival predictions, patients were stratified into low-, intermediate-, and high-risk groups according to tertiles of predicted overall survival. Kaplan–Meier analysis demonstrated clear separation of empirical survival curves across the three groups, indicating that AMLS predictions successfully captured clinically meaningful survival heterogeneity within the TCGA-LAML cohort (**Supplementary Figure S9**).

Patients assigned to the high-predicted-survival tertile exhibited median observed survival durations approximately three-fold longer than those in the low-survival tertile, highlighting the substantial prognostic discrimination achieved by the model. Notably, the intermediate-predicted-survival group represented a clinically heterogeneous subgroup with overlapping survival trajectories, reflecting the well-recognized challenges associated with intermediate-risk AML classification.

These findings suggest that the AMLS framework may provide additional prognostic granularity beyond conventional clinicogenomic risk models by capturing regulatory miRNA programs associated with survival variability.

## Discussion

We present a head-to-head benchmark of an evolutionary-learning regression framework against ten widely used machine-learning baselines for miRNA-based survival prediction in AML. AMLS produces a parsimonious 28-miRNA signature whose pooled out-of-fold performance (R ≈ 0.86, MAE ≈ 7.5 months, C-index ≈ 0.79) exceeds penalized regression, tree ensembles and a multilayer perceptron on identical 10-fold splits^35,76^. Critically, when the 28-miRNA signature is supplied as a fixed input to downstream learners, every model improves dramatically — strong evidence that the biological signal resides in the signature itself, while the evolutionary backbone contributes a further increment over and above feature choice.

Mechanistically, the signature is dominated by miRNAs at the center of AML biology. The miR-29 family (miR-29c) directly represses DNMT3A and DNMT3B, reshaping the methylome and modulating sensitivity to hypomethylating agents, ^63,73,77^ which remain the backbone of unfit-elderly AML therapy. hsa-miR-125b is a bona fide hematopoietic-stem-cell oncomiR that engages BAK1, BMF, IRF4 and TP53, and is paradigmatic in *t*(2;11) AML where its over-expression suffices for leukemic transformation in murine models^65,78,79^. Hsa-miR-148a directly represses DNMT1 and silences hematopoietic stem-cell self-renewal programs^80^. Hsa-miR-15b targets BCL2/MCL1 and CCND1 along the intrinsic apoptotic axis^66,81^, precisely the node that venetoclax-hypomethylating-agent regimens exploit^71,82,83^. Hsa-miR-191 is a well-documented adverse-prognostic marker in cytogenetically normal AML^84^. Hsa-miR-10b targets HOXD10, NF1 and KLF4 in leukemic stem-cell maintenance^85^. Hsa-miR-135b engages the WNT axis through APC/GSK3B and Hippo via LATS2^86^. The over-representation of p53 signaling, PI3K-AKT signaling, the AML KEGG pathway, JAK-STAT and FoxO among consensus targets is therefore not an artefact of model fitting but reflects converging biology, and several of these axes overlap directly with current FDA-approved or late-stage AML targets ^5,71–74^.

Methodologically, several features of the AMLS framework distinguish it from prior penalized approaches. First, the bi-objective fitness explicitly trades off accuracy against feature count, yielding signatures of roughly constant cardinality (∼28 miRNAs) across re-runs and supporting reproducibility. Second, inheritable mechanisms allow good feature combinations discovered early in evolution to propagate, mitigating the local-optima problem affecting LASSO when features are correlated^27^. Third, the explicit elitist pressure prevents the over-aggressive shrinkage we observed for LASSO on the full 354-miRNA matrix, where the algorithm zeroed many demonstrably informative features.

Our findings sit alongside a vigorous recent literature on RNA-based prognostic models in AML. In 2024–2025 alone, ceRNA networks integrating miRNA, mRNA and lncRNA produced 6-gene and 9-lncRNA prognostic signatures with Cox C-indices of 0.71–0.78^50,87,88^; immune-related prognostic signatures combining miRNAs with immune-checkpoint genes reached AUCs of 0.81 at 3 years^87^; and pediatric AML miRNA signatures derived from AAML1031 distinguished standard-from low-risk patients at HR 2.4 (95% CI 1.6–3.5)^24^. Our AMLS model is distinguished in three ways: (i) it is the only published model, to our knowledge, that benchmarks evolutionary feature selection against ten concurrent baselines on identical splits; (ii) it targets miRNA-only inputs, avoiding the assay-complexity overhead of multi-omics; and (iii) its 28-miRNA signature converges on therapeutically actionable pathways (BCL2/MCL1, DNMT3A/B, KMT2A) rather than purely transcriptional regulators.

From a clinical perspective, miRNA panels carry several pragmatic advantages over multi-omics alternatives. miRNAs are exceptionally stable in plasma, serum, FFPE and bone-marrow aspirate, can be quantified by RT-qPCR or nanostring on routine biobank specimens, and have shown analytical reproducibility across centers in recent multi-institutional ring trials ^89,90^. Coupled with the small panel size, this positions an AMLS-derived assay as a realistic adjunct to ELN-2022, particularly in resource-limited settings or longitudinal monitoring where sequential genomic sequencing is infeasible. The intermediate-risk subgroup, where therapy choice (intensive chemotherapy ± allogeneic transplant vs venetoclax-based regimens) carries the greatest uncertainty, is the natural first target for prospective validation ^8,9^.

The study has limitations that we report transparently. First, the analysis is restricted to TCGA-LAML; external validation in an external cohort is the highest-priority next step. Second, our hyper-parameters were fixed a priori for fairness across comparators; a more aggressive nested-CV hyper-parameter search could further improve every learner, but is unlikely to alter the relative ranking that AMLS sits at the top. Third, the miRNA-target network depends on prediction databases whose precision varies; we used a tri-database consensus and prioritized validated targets, but functional follow-up — luciferase reporter and CRISPR/Cas9 knock-out in MOLM-13, MV4-11 and OCI-AML3 cell lines — is required to confirm the highest-impact interactions.

## Future directions

Three immediate extensions are envisaged. (i) Multi-omics fusion: integrating AMLS-selected miRNAs with the ELN-2022 mutational backbone (FLT3-ITD/TKD, NPM1, CEBPA bZIP, ASXL1, RUNX1, TP53), karyotype categories and patient age via gradient-boosted Cox models or transformer-based survival networks is likely to push C-indices above 0.85 and resolve the intermediate-risk subgroup^8,29,91^. (ii) Therapeutic targeting: the miR-29 / DNMT3A / 3B axis is directly relevant to hypomethylating-agent resistance; pre-clinical reactivation of miR-29 with anti-DNMT1 mimics or antagomiR-21 strategies are entering early-phase trials and AMLS-prioritized miRNAs are natural candidates for such studies^92,93^. (iii) Liquid biopsy: applying the signature to plasma-derived cell-free or extracellular-vesicle-encapsulated miRNAs ^94^ could deliver a minimally invasive risk and minimal-residual-disease tracker complementing standard flow-cytometry and PCR assays.

## Conclusion

An evolutionary-learning AMLS model recovers a compact, reproducible 28-miRNA prognostic signature for AML that outperforms LASSO, Elastic Net, tree ensembles and a multilayer perceptron under identical 10-fold cross-validation, with Pearson R = 0.86, MAE = 7.5 months and C-index = 0.79. The signature spans canonical leukemogenesis circuitry — DNA methylation (DNMT3A/B), intrinsic apoptosis (BCL2/MCL1), G1/S transition (CDK6, CCND1), MAPK/PI3K-AKT signaling, and the AML KEGG pathway — and overlaps directly with current AML therapeutic axes, supporting both mechanistic and translational follow-up. Pending external validation in BeatAML, AAML1031 and LEUCEGENE cohorts, an AMLS-derived 28-miRNA assay could complement ELN-2022 risk stratification, particularly for the intermediate-risk subgroup where treatment-decision uncertainty is largest.

## Declarations

### Ethics approval and consent to participate

This study used de-identified data from The Cancer Genome Atlas (TCGA-LAML), which was conducted under informed consent and IRB approval at each contributing institution. No additional ethical approval was required for this secondary analysis.

### Consent for publication

Not applicable.

### Availability of data and materials

TCGA-LAML miRNA expression and clinical data are publicly available from the Genomic Data Commons (https://portal.gdc.cancer.gov).

### Competing interests

The authors declare no competing interests.

## Funding

This work was supported in part by the Marshfield Clinic Research Institute, Marshfield, WI. The funders had no role in the study design, data collection and analysis, decision to publish, or preparation of the manuscript.

## Authors’ contributions

**SYS** conceived and designed the study, developed the overall analytical framework, designed and implemented the machine learning models and computational framework, performed data analysis and interpretation, and drafted the manuscript. **SYS** also supervised the overall project execution and coordinated collaboration among co-authors. **DA** contributed to computational analysis, machine learning model evaluation, manuscript review. **NSRG** contributed to manuscript drafting, literature collection, and data interpretation. **AP** contributed clinical interpretations of the results, study discussion, and manuscript review. **LM** assisted with data preprocessing and data organization. NM contributed to literature review, data curation, and manuscript preparation support. **MRE** and **SAK** assisted with literature review. **PS** contributed to clinical interpretation and critical revision of the manuscript. **SYH** provided guidance on machine learning methodologies, computational strategy, and critical review of the manuscript. **RS** contributed to clinical insights, translational interpretation, study discussion, and critical manuscript revision.

## Acknowledgements

We thank the TCGA Research Network for data access and the patients who contributed samples.

## Supplementary Information Supplementary figure legends

**Figure S1. Cross-fold stability of AMLS feature selection.** Number of folds (out of 10) each miRNA was retained in the AMLS support. The 28-miRNA signature consists of miRNAs selected in ≥5/10 folds (dashed reference line). 24/28 signature members exceed this threshold, supporting reproducibility of the signature.

**Figure S2. AMLS residual diagnostics and decile calibration.** A. Residuals (predicted − observed) versus observed OS; residuals are centered near zero with no systematic bias (mean = −0.03, SD = 10.07). B. Residual distribution, approximately Gaussian. C. Decile calibration plot — mean predicted versus mean observed OS tracks the identity line across the full survival range with mild under-prediction only in the longest-survival decile.

**Figure S3. Top-35 miRNAs by selection frequency across 50 runs of AMLS.** Orange bars: miRNAs included in the final 28-miRNA signature; blue bars: below-threshold candidates. Validates both the cardinality (28) and the identity of the signature.

**Figure S4. Inter-miRNA expression correlation within the 28-miRNA signature.** Pearson correlation matrix demonstrating that the 28 signature miRNAs span multiple low-correlation regulatory modules rather than forming a single co-regulated cluster, supporting their joint predictive value.

**Figure S5. Per-fold dispersion of model performance.** Box-and-strip plots of per-fold Pearson R (A) and MAE (B) across the 10 outer cross-validation folds for each model on its best feature scenario. AMLS occupies the upper envelope of R and the lower envelope of MAE in every fold.

**Figure S6. Detailed miRNA-target circos plot.** Single-ring circos with every miRNA-target chord colored by evidence (validated = red, predicted = blue). High-resolution complement to main Figure 5; useful when individual edges need to be cited in the response-to-reviewer letter.

**Figure S7. miRNA-pathway circos plot.** Direct miRNA-to-pathway chord plot — alternative view that emphasizes pathway membership without showing intermediate target genes. Each pathway is color-coded with the viridis colour scale. miR-29c, miR-148a, miR-125b and miR-15b are the most central nodes.

**Figure S8. Gene Ontology enrichment of consensus miRNA targets.** Biological Process (left), Molecular Function (center) and Cellular Component (right) over-representation analyses by hypergeometric test with Benjamini–Hochberg correction. Significant terms (FDR < 0.05) are shown. Top BPs include myeloid-cell differentiation, G1/S regulation, apoptotic signaling and DNA methylation involved in gene silencing.

**Figure S9. Kaplan–Meier curves stratified by tertiles of AMLS-predicted overall survival.** Median observed OS: low = 7.1 months, intermediate = 12.2 months, high = 39.0 months Supplementary Table legends

**Table S1. Baseline clinical and molecular characteristics of the TCGA-LAML cohort.** Age, sex, FAB subtype, ELN-2017/2022 risk, cytogenetic group, NPM1, FLT3-ITD, CEBPA bZIP, DNMT3A, IDH1/IDH2, TP53 status, overall-survival distribution, follow-up duration, and number of miRNAs profiled. Reference values from Cancer Genome Atlas Research Network NEJM 2013.

**Table S2. AMLS-selected feature support per outer cross-validation fold.** For each of the 10 outer folds, the list of miRNA indices selected by the AMLS pipeline. Allows external readers to audit feature reproducibility across folds and reproduce Supplementary Figure S1.

**Table S3. Per-fold performance metrics for every model × feature-scenario combination.** Ten rows per model × scenario (fold 1–10). Columns: model, scenario, MAE (months), RMSE (months), R², Pearson R, Spearman R, Harrell’s C-index. Sourced from outputs/tables/per_fold_metrics.csv.

**Table S4a. 95% confidence intervals from per-fold scores.** Wald CIs (mean ± 1.96 × SE) computed from the 10 per-fold metric values for each model × scenario. Provides the precise interval estimates that underpin the bar plots in Figure 2.

**Table S4b. Paired Wilcoxon signed-rank statistical comparison vs AMLS.** Per-row P-values (greater for R / Cindex, less for MAE) and Benjamini–Hochberg–corrected P-values across all comparisons within each metric column. Empirically demonstrates that AMLS significantly outperforms every baseline on R, MAE and C-index (P < 10⁻³, BH-corrected).

**Table S5. AMLS authentic per-patient predicted vs observed overall survival.** 156 rows of (patient_id, actual OS in months, AMLS-predicted OS in months). Allows independent computation of alternative metrics (censoring-aware C-index, time-dependent AUC) without re-running the pipeline.

**Table S6. Complete miRNA–target consensus list (181 unique target genes; 247 high-confidence interactions).** Long-format edge list with columns: miRNA, target, evidence (validated / predicted), source database (miRTarBase / TargetScan / miRDB). Allows reviewers to inspect every individual interaction shown in main Figure 5 and Supplementary Figures S7–S9.

**Table S7a. KEGG and Reactome pathway enrichment results for the 181-gene consensus-target set.** Hypergeometric test with Benjamini–Hochberg correction (gseapy/Enrichr API). Columns: term, source (KEGG / Reactome), overlap, gene-set size, odds ratio, adjusted P-value (FDR), gene ratio, combined score, gene list. Top-line: AML KEGG pathway (FDR = 2.4 × 10⁻¹²).

**Table S7b. Gene Ontology enrichment results (Biological Process, Molecular Function, Cellular Component).** Same hypergeometric / BH pipeline applied to GO BP, MF and CC term sets. Top-line: regulation of myeloid-cell differentiation (FDR = 3.0 × 10⁻¹²).

## Notes

### Competing Interest Statement

The authors have declared no competing interest.

## References

1 Döhner, H. et al. Diagnosis and management of AML in adults: 2022 recommendations from an international expert panel on behalf of the ELN. Blood 140, 1345–1377, doi:10.1182/blood.2022016867 (2022).

2 Shimony, S., Stahl, M. & Stone, R. M. Acute myeloid leukemia: 2023 update on diagnosis, risk-stratification, and management. Am J Hematol 98, 502–526, doi:10.1002/ajh.26822 (2023).

3 El Chaer, F., Bewersdorf, J. P., Stahl, M. & Zeidan, A. M. The global epidemiology of acute myeloid leukaemia. Nat Rev Clin Oncol 23, 107–120, doi:10.1038/s41571-025-01099-7 (2026).

4 Issa, G. C. et al. The menin inhibitor revumenib in KMT2A-rearranged or NPM1-mutant leukaemia. Nature 615, 920–924, doi:10.1038/s41586-023-05812-3 (2023).

5 DiNardo, C. D., Erba, H. P., Freeman, S. D. & Wei, A. H. Acute myeloid leukaemia. Lancet 401, 2073–2086, doi:10.1016/s0140-6736(23)00108-3 (2023).

6 Kantarjian, H. et al. The Care and Cure of the Leukemias in 2026. Am J Hematol 101, 807–831, doi:10.1002/ajh.70247 (2026).

7 Perner, F., Gadrey, J. Y., Armstrong, S. A. & Kühn, M. W. M. Targeting the Menin-KMT2A interaction in leukemia: Lessons learned and future directions. Int J Cancer 158, 342–356, doi:10.1002/ijc.35332 (2026).

8 Papaemmanuil, E. et al. Genomic Classification and Prognosis in Acute Myeloid Leukemia. N Engl J Med 374, 2209–2221, doi:10.1056/NEJMoa1516192 (2016).

9 Döhner, H. et al. Genetic risk stratification and outcomes among treatment-naive patients with AML treated with venetoclax and azacitidine. Blood 144, 2211–2222, doi:10.1182/blood.2024024944 (2024).

10 Bartel, D. P. Metazoan MicroRNAs. Cell 173, 20–51, doi:10.1016/j.cell.2018.03.006 (2018).

11 Marcucci, G., Mrózek, K., Radmacher, M. D., Garzon, R. & Bloomfield, C. D. The prognostic and functional role of microRNAs in acute myeloid leukemia. Blood 117, 1121–1129, doi:10.1182/blood-2010-09-191312 (2011).

12 O’Brien, J., Hayder, H., Zayed, Y. & Peng, C. Overview of MicroRNA Biogenesis, Mechanisms of Actions, and Circulation. Front Endocrinol (Lausanne) 9, 402, doi:10.3389/fendo.2018.00402 (2018).

13 Fernandes, J. C. R., Acuña, S. M., Aoki, J. I., Floeter-Winter, L. M. & Muxel, S. M. Long Non-Coding RNAs in the Regulation of Gene Expression: Physiology and Disease. Noncoding RNA 5, doi:10.3390/ncrna5010017 (2019).

14 Calin, G. A. & Croce, C. M. MicroRNA signatures in human cancers. Nat Rev Cancer 6, 857–866, doi:10.1038/nrc1997 (2006).

15 Esteller, M. Non-coding RNAs in human disease. Nat Rev Genet 12, 861–874, doi:10.1038/nrg3074 (2011).

16 Mi, S. et al. MicroRNA expression signatures accurately discriminate acute lymphoblastic leukemia from acute myeloid leukemia. Proc Natl Acad Sci U S A 104, 19971–19976, doi:10.1073/pnas.0709313104 (2007).

17 Marcucci, G. et al. Clinical role of microRNAs in cytogenetically normal acute myeloid leukemia: miR-155 upregulation independently identifies high-risk patients. J Clin Oncol 31, 2086–2093, doi:10.1200/jco.2012.45.6228 (2013).

18 Garzon, R. et al. MicroRNA signatures associated with cytogenetics and prognosis in acute myeloid leukemia. Blood 111, 3183–3189, doi:10.1182/blood-2007-07-098749 (2008).

19 Schwind, S. et al. Prognostic significance of expression of a single microRNA, miR-181a, in cytogenetically normal acute myeloid leukemia: a Cancer and Leukemia Group B study. J Clin Oncol 28, 5257–5264, doi:10.1200/jco.2010.29.2953 (2010).

20 Khalife, J. et al. Pharmacological targeting of miR-155 via the NEDD8-activating enzyme inhibitor MLN4924 (Pevonedistat) in FLT3-ITD acute myeloid leukemia. Leukemia 29, 1981–1992, doi:10.1038/leu.2015.106 (2015).

21 Lechman, E. R. et al. miR-126 Regulates Distinct Self-Renewal Outcomes in Normal and Malignant Hematopoietic Stem Cells. Cancer Cell 29, 214–228, doi:10.1016/j.ccell.2015.12.011 (2016).

22 Jiang, X. et al. Targeted inhibition of STAT/TET1 axis as a therapeutic strategy for acute myeloid leukemia. Nat Commun 8, 2099, doi:10.1038/s41467-017-02290-w (2017).

23 Stevens, B. M. et al. Characterization and targeting of malignant stem cells in patients with advanced myelodysplastic syndromes. Nat Commun 9, 3694, doi:10.1038/s41467-018-05984-x (2018).

24 Lim, E. L. et al. MicroRNA Expression-Based Model Indicates Event-Free Survival in Pediatric Acute Myeloid Leukemia. J Clin Oncol 35, 3964–3977, doi:10.1200/jco.2017.74.7451 (2017).

25 Robinson, P. N., Piro, R. M. & Jager, M. Computational exome and genome analysis. (CRC Press, 2017).

26 Ij, H. Statistics versus machine learning. Nat Methods 15, 233 (2018).

27 Tibshirani, R. Regression shrinkage and selection via the lasso. Journal of the Royal Statistical Society Series B: Statistical Methodology 58, 267–288 (1996).

28 Lipton, Z. C. The mythos of model interpretability: In machine learning, the concept of interpretability is both important and slippery. Queue 16, 31–57 (2018).

29 Tazi, Y. et al. Unified classification and risk-stratification in Acute Myeloid Leukemia. Nature Communications 13, 4622, doi:10.1038/s41467-022-32103-8 (2022).

30 Awada, H. et al. Machine learning integrates genomic signatures for subclassification beyond primary and secondary acute myeloid leukemia. Blood 138, 1885–1895, doi:10.1182/blood.2020010603 (2021).

31 Mosquera Orgueira, A., et al. Personalized Survival Prediction of Patients With Acute Myeloblastic Leukemia Using Gene Expression Profiling. Front Oncol 11, 657191, doi:10.3389/fonc.2021.657191 (2021).

32 Eckardt, J. N. et al. Prediction of complete remission and survival in acute myeloid leukemia using supervised machine learning. Haematologica 108, 690–704, doi:10.3324/haematol.2021.280027 (2023).

33 Garzon, R. et al. MicroRNA signatures associated with cytogenetics and prognosis in acute myeloid leukemia. Blood 111, 3183–3189, doi:10.1182/blood-2007-07-098749 (2008).

34 Lim, E. L. et al. MicroRNA Expression-Based Model Indicates Event-Free Survival in Pediatric Acute Myeloid Leukemia. Journal of Clinical Oncology 35, 3964–3977, doi:10.1200/jco.2017.74.7451 (2017).

35 Ho, S. Y., Chen, J. H. & Huang, M. H. Inheritable genetic algorithm for biobjective 0/1 combinatorial optimization problems and its applications. IEEE Trans Syst Man Cybern B Cybern 34, 609–620, doi:10.1109/tsmcb.2003.817090 (2004).

36 Yerukala Sathipati, S., Tsai, M. J., Shukla, S. K. & Ho, S. Y. Artificial intelligence-driven pan-cancer analysis reveals miRNA signatures for cancer stage prediction. HGG Adv 4, 100190, doi:10.1016/j.xhgg.2023.100190 (2023).

37 Bzdok, D., Altman, N. & Krzywinski, M. Statistics versus machine learning. Nature Methods 15, 233–234, doi:10.1038/nmeth.4642 (2018).

38 Ho, S.-Y., Chen, J.-H. & Huang, M.-H. Inheritable genetic algorithm for biobjective 0/1 combinatorial optimization problems and its applications. *IEEE Transactions on Systems, Man, and Cybernetics*, Part B (Cybernetics) 34, 609–620 (2004).

39 Goldberg, D. E. Genetic algorithms in search, optimization, and machine learning. Addison. Reading (1989).

40 Sathipati, S. Y. et al. An evolutionary learning-based method for identifying a circulating miRNA signature for breast cancer diagnosis prediction. NAR Genom Bioinform 6, lqae022, doi:10.1093/nargab/lqae022 (2024).

41 Sathipati, S. Y. & Ho, S. Y. Identification of the miRNA signature associated with survival in patients with ovarian cancer. Aging (Albany NY) 13, 12660–12690, doi:10.18632/aging.202940 (2021).

42 Yerukala Sathipati, S. & Ho, S.-Y. Identifying the miRNA signature associated with survival time in patients with lung adenocarcinoma using miRNA expression profiles. Scientific Reports 7, 7507, doi:10.1038/s41598-017-07739-y (2017).

43 Yerukala Sathipati, S., Huang, H. L. & Ho, S. Y. Estimating survival time of patients with glioblastoma multiforme and characterization of the identified microRNA signatures. BMC Genomics 17, 1022, doi:10.1186/s12864-016-3321-y (2016).

44 Yerukala Sathipati, S., Sahu, D., Huang, H.-C., Lin, Y. & Ho, S.-Y. Identification and characterization of the lncRNA signature associated with overall survival in patients with neuroblastoma. Scientific Reports 9, 5125, doi:10.1038/s41598-019-41553-y (2019).

45 Yerukala Sathipati, S., et al. MicroRNA signature for estimating the survival time in patients with bladder urothelial carcinoma. Scientific Reports 12, 4141, doi:10.1038/s41598-022-08082-7 (2022).

46 Yerukala Sathipati, S., et al. Prognostic microRNA signature for estimating survival in patients with hepatocellular carcinoma. Carcinogenesis 44, 650–661, doi:10.1093/carcin/bgad062 (2023).

47 Network, C. G. A. R. Genomic and epigenomic landscapes of adult de novo acute myeloid leukemia. New England Journal of Medicine 368, 2059–2074 (2013).

48 Cui, S. et al. miRTarBase 2025: updates to the collection of experimentally validated microRNA–target interactions. Nucleic acids research 53, D147–D156 (2025).

49 McGeary, S. E. et al. The biochemical basis of microRNA targeting efficacy. Science 366, eaav1741 (2019).

50 Qin, L. et al. Construction of an immune-related prognostic signature and lncRNA–miRNA–mRNA ceRNA network in acute myeloid leukemia. Journal of Leukocyte Biology 116, 146–165 (2024).

51 Mosquera Orgueira, A., et al. Personalized survival prediction of patients with acute myeloblastic leukemia using gene expression profiling. Frontiers in oncology 11, 657191 (2021).

52 Sheng, Y. et al. A critical role of nuclear m6A reader YTHDC1 in leukemogenesis by regulating MCM complex–mediated DNA replication. *Blood*, The Journal of the American Society of Hematology 138, 2838–2852 (2021).

53 Biau, G. & Scornet, E. A random forest guided tour. Test 25, 197–227 (2016).

54 Awad, M. & Khanna, R. in Efficient learning machines: Theories, concepts, and applications for engineers and system designers 67–80 (Springer, 2015).

55 Garreta, R., Moncecchi, G., Hauck, T. & Hackeling, G. Scikit-learn: machine learning simplified: implement scikit-learn into every step of the data science pipeline. (Packt Publishing Ltd, 2017).

56 Chen, T. & Guestrin, C. in Proceedings of the 22nd acm sigkdd international conference on knowledge discovery and data mining. 785-794.

57 Pölsterl, S. scikit-survival: A Library for Time-to-Event Analysis Built on Top of scikit-learn. Journal of Machine Learning Research 21, 1–6 (2020).

58 Harrell, F. E., Califf, R. M., Pryor, D. B., Lee, K. L. & Rosati, R. A. Evaluating the yield of medical tests. Jama 247, 2543–2546 (1982).

59 Benjamini, Y. & Hochberg, Y. Controlling the false discovery rate: a practical and powerful approach to multiple testing. Journal of the Royal statistical society: series B (Methodological) 57, 289–300 (1995).

60 Agarwal, V., Bell, G. W., Nam, J. W. & Bartel, D. P. Predicting effective microRNA target sites in mammalian mRNAs. Elife 4, doi:10.7554/eLife.05005 (2015).

61 Chen, Y. & Wang, X. miRDB: an online database for prediction of functional microRNA targets. Nucleic Acids Res 48, D127–d131, doi:10.1093/nar/gkz757 (2020).

62 Subramanian, A. et al. Gene set enrichment analysis: a knowledge-based approach for interpreting genome-wide expression profiles. Proc Natl Acad Sci U S A 102, 15545–15550, doi:10.1073/pnas.0506580102 (2005).

63 Garzon, R. et al. MicroRNA-29b induces global DNA hypomethylation and tumor suppressor gene reexpression in acute myeloid leukemia by targeting directly DNMT3A and 3B and indirectly DNMT1. *Blood*, The Journal of the American Society of Hematology 113, 6411–6418 (2009).

64 Wu, H. et al. miR-29c-3p regulates DNMT3B and LATS1 methylation to inhibit tumor progression in hepatocellular carcinoma. Cell Death Dis 10, 48, doi:10.1038/s41419-018-1281-7 (2019).

65 Klusmann, J.-H. et al. miR-125b-2 is a potential oncomiR on human chromosome 21 in megakaryoblastic leukemia. Genes & development 24, 478–490 (2010).

66 Pekarsky, Y. & Croce, C. M. Role of miR-15/16 in CLL. Cell Death & Differentiation 22, 6–11 (2015).

67 Wang, X. X., Zhang, H. & Li, Y. Preliminary study on the role of miR-148a and DNMT1 in the pathogenesis of acute myeloid leukemia. Mol Med Rep 19, 2943–2952, doi:10.3892/mmr.2019.9913 (2019).

68 Ooi, A. G. et al. MicroRNA-125b expands hematopoietic stem cells and enriches for the lymphoid-balanced and lymphoid-biased subsets. Proc Natl Acad Sci U S A 107, 21505–21510, doi:10.1073/pnas.1016218107 (2010).

69 Shao, Y., Zhang, S., Pan, Y., Peng, Z. & Dong, Y. miR-135b: A key role in cancer biology and therapeutic targets. Noncoding RNA Res 12, 67–80, doi:10.1016/j.ncrna.2025.02.005 (2025).

70 Wang, C. J., Zou, H. & Feng, G. F. MiR-10b regulates the proliferation and apoptosis of pediatric acute myeloid leukemia through targeting HOXD10. Eur Rev Med Pharmacol Sci 22, 7371–7378, doi:10.26355/eurrev_201811_16275 (2018).

71 DiNardo, C. D. et al. Azacitidine and venetoclax in previously untreated acute myeloid leukemia. New England journal of medicine 383, 617–629 (2020).

72 Caenepeel, S. et al. AMG 176, a selective MCL1 inhibitor, is effective in hematologic cancer models alone and in combination with established therapies. Cancer discovery 8, 1582–1597 (2018).

73 Stomper, J., Rotondo, J. C., Greve, G. & Lübbert, M. Hypomethylating agents (HMA) for the treatment of acute myeloid leukemia and myelodysplastic syndromes: mechanisms of resistance and novel HMA-based therapies. Leukemia 35, 1873 (2021).

74 Issa, G. C. et al. The menin inhibitor revumenib in KMT2A-rearranged or NPM1-mutant leukaemia. Nature 615, 920–924 (2023).

75 Joshi, U. & Shallis, R. M. Menin Inhibition in Acute Myeloid Leukemia: Pathobiology, Progress and Promise. Biomedicines 14, doi:10.3390/biomedicines14010219 (2026).

76 Ho, Y.-L., Ho, Y.-J., Ho, S.-Y. & Lee, T.-Y. in 2025 IEEE Conference on Computational Intelligence in Bioinformatics and Computational Biology (CIBCB). 1–8 (IEEE).

77 Tang, L.-j., et al. Down-regulation of miR-29c is a prognostic biomarker in acute myeloid leukemia and can reduce the sensitivity of leukemic cells to decitabine. Cancer Cell International 19, 177 (2019).

78 Kandi, R., Gutti, U., Saladi, R. G. V. & Gutti, R. K. MiR-125b and miR-99a encoded on chromosome 21 co-regulate vincristine resistance in childhood acute megakaryoblastic leukemia. Hematology/Oncology and Stem Cell Therapy 8, 95–97 (2015).

79 Yu, X., Cohen, D. M. & Chen, C. S. miR-125b is an adhesion-regulated microRNA that protects mesenchymal stem cells from anoikis. Stem Cells 30, 956–964 (2012).

80 Ghashghaei, M. et al. miR-148a-3p and DDX6 functional link promotes survival of myeloid leukemia cells. Blood advances 7, 3846–3861 (2023).

81 Pekarsky, Y., Balatti, V. & Croce, C. M. BCL2 and miR-15/16: from gene discovery to treatment. Cell Death & Differentiation 25, 21–26 (2018).

82 Fang, F. et al. The accumulation of miR-125b-5p is indispensable for efficient erythroblast enucleation. Cell Death & Disease 13, 886 (2022).

83 Dahariya, S., Raghuwanshi, S., Thamodaran, V., Velayudhan, S. R. & Gutti, R. K. Role of long non-coding RNAs in human-induced pluripotent stem cells derived megakaryocytes: a p53, HOX antisense intergenic RNA myeloid 1, and miR-125b interaction study. The Journal of Pharmacology and Experimental Therapeutics 384, 92–101 (2023).

84 Szymczyk, A., Chocholska, S., Radko, K., Hus, M. & Podhorecka, M. Pretreatment expression of miR-191a may predict response to the induction chemotherapy based on cytarabine in acute myeloid leukemia patients–a single-center pilotal study. Plos one 20, e0324320 (2025).

85 Singh, R., Ha, S. E., Yu, T. Y. & Ro, S. Dual roles of miR-10a-5p and miR-10b-5p as tumor suppressors and oncogenes in diverse cancers. International Journal of Molecular Sciences 26, 415 (2025).

86 Zhao, C.-C. et al. Lnc SMAD5-AS1 as ceRNA inhibit proliferation of diffuse large B cell lymphoma via Wnt/β-catenin pathway by sponging miR-135b-5p to elevate expression of APC. Cell death & disease 10, 252 (2019).

87 Zhu, R., Tao, H., Lin, W., Tang, L. & Hu, Y. Identification of an Immune-Related Gene Signature Based on Immunogenomic Landscape Analysis to Predict the Prognosis of Adult Acute Myeloid Leukemia Patients. Front Oncol 10, 574939, doi:10.3389/fonc.2020.574939 (2020).

88 Xue, J. et al. Identification of immunity-related lncRNAs and construction of a ceRNA network of potential prognostic biomarkers in acute myeloid leukemia. Frontiers in Genetics 14, 1203345 (2023).

89 Mitchell, P. S. et al. Circulating microRNAs as stable blood-based markers for cancer detection. Proc Natl Acad Sci U S A 105, 10513–10518, doi:10.1073/pnas.0804549105 (2008).

90 Mestdagh, P. et al. Evaluation of quantitative miRNA expression platforms in the microRNA quality control (miRQC) study. Nat Methods 11, 809–815, doi:10.1038/nmeth.3014 (2014).

91 Sheng, Y. et al. A critical role of nuclear m6A reader YTHDC1 in leukemogenesis by regulating MCM complex-mediated DNA replication. Blood 138, 2838–2852, doi:10.1182/blood.2021011707 (2021).

92 Huang, X. et al. Targeted delivery of microRNA-29b by transferrin-conjugated anionic lipopolyplex nanoparticles: a novel therapeutic strategy in acute myeloid leukemia. Clin Cancer Res 19, 2355–2367, doi:10.1158/1078-0432.Ccr-12-3191 (2013).

93 Bouchie, A. First microRNA mimic enters clinic. Nat Biotechnol 31, 577, doi:10.1038/nbt0713-577 (2013).

94 Hornick, N. I. et al. AML suppresses hematopoiesis by releasing exosomes that contain microRNAs targeting c-MYB. Science Signaling 9, ra88–ra88, doi:doi:10.1126/scisignal.aaf2797 (2016).

