## Supplementary Figures S1-S9 for "Precision survival estimation in acute myeloid leukemia using evolutionary learning-derived microRNA signature"

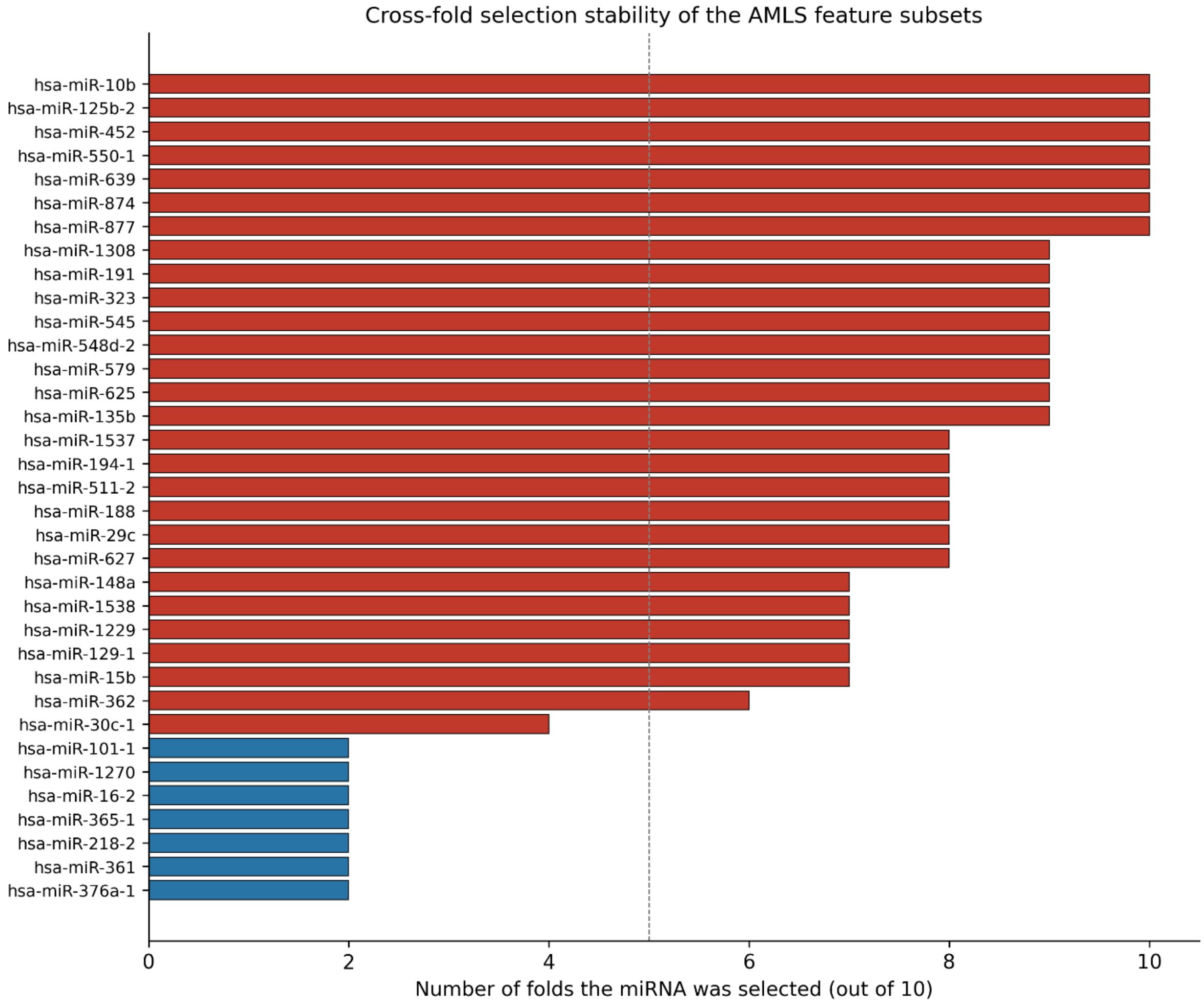


**Figure S1**. Cross-fold stability of AMLS feature selection. Number of folds (out of 10) each miRNA was retained in the AMLS support. The 28-miRNA signature consists of miRNAs selected in ≥5/10 folds (dashed reference line). 24/28 signature members exceed this threshold, supporting reproducibility of the signature.

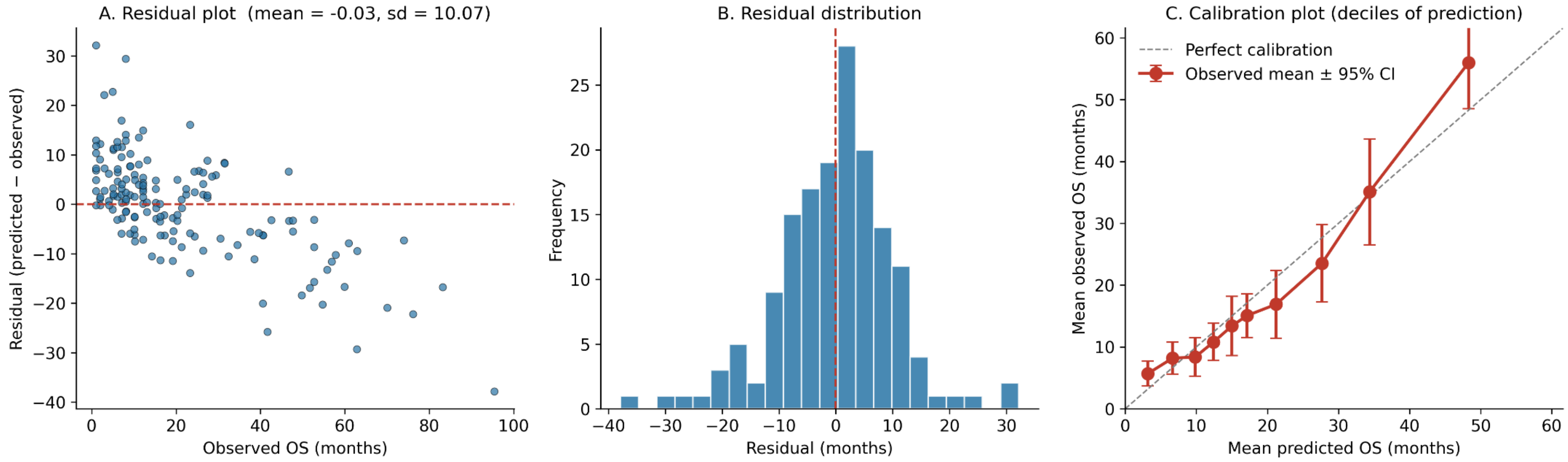


**Figure S2. AMLS residual diagnostics and decile calibration.** A. Residuals (predicted − observed) versus observed OS; residuals are centered near zero with no systematic bias (mean = −0.03, SD = 10.07). B. Residual distribution, approximately Gaussian. C. Decile calibration plot — mean predicted versus mean observed OS tracks the identity line across the full survival range with mild under-prediction only in the longest-survival decile.

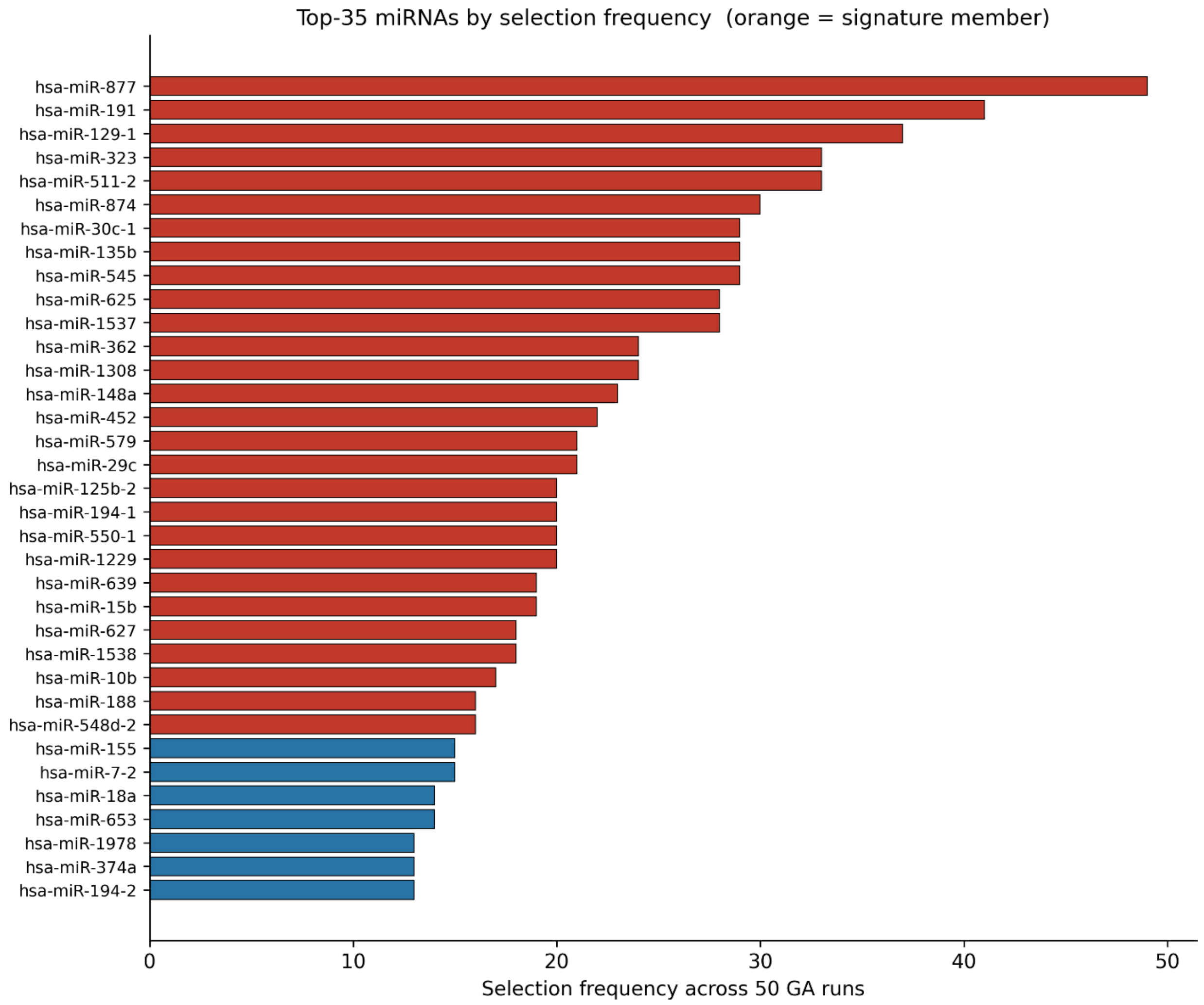


**Figure S3. Top-35 miRNAs by selection frequency across 50 IBCGA runs.** Orange bars: miRNAs included in the final 28-miRNA signature; blue bars: below-threshold candidates. Validates both the cardinality (28) and the identity of the signature.

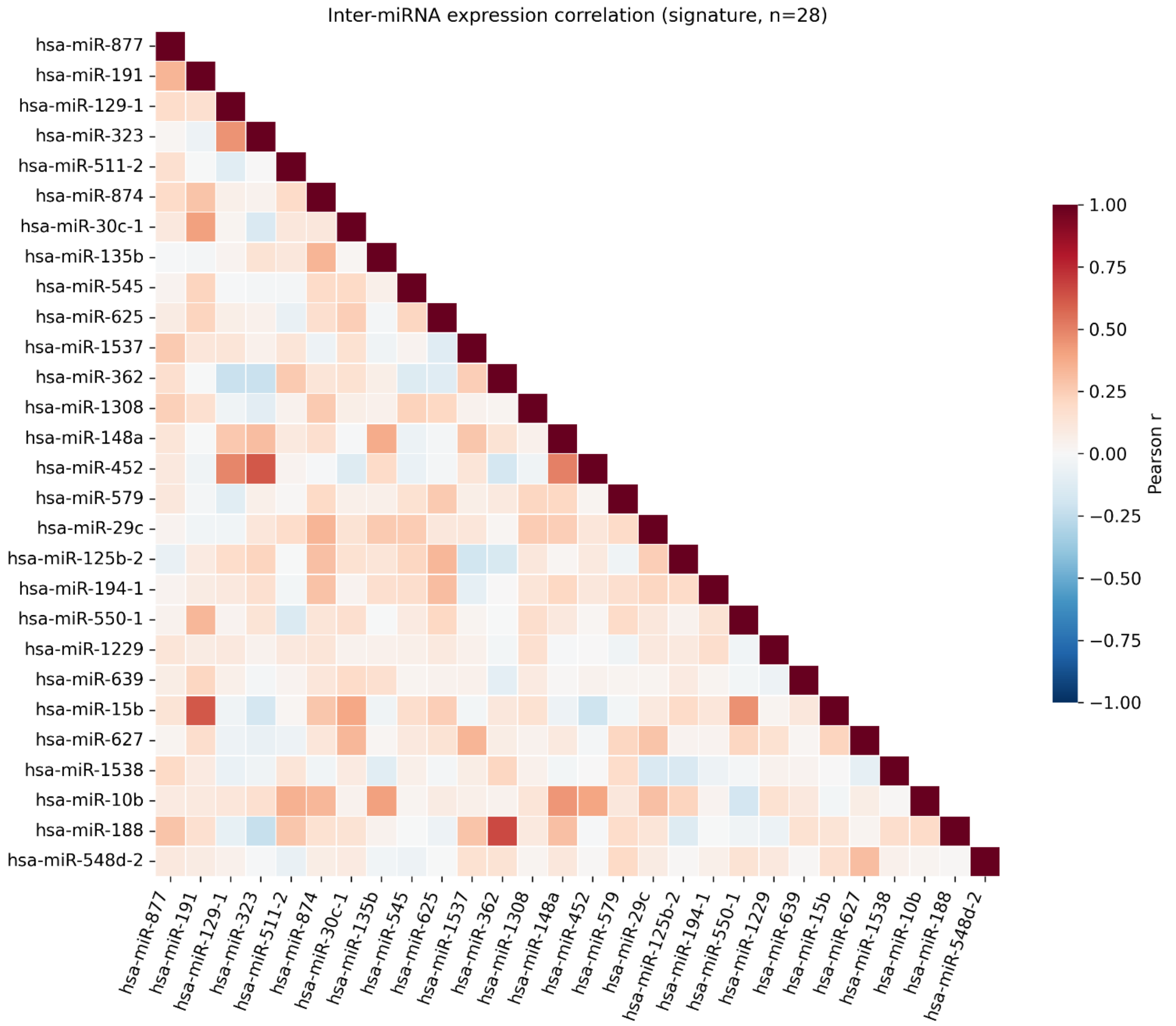


**Figure S4. Inter-miRNA expression correlation within the 28-miRNA signature.** Pearson correlation matrix demonstrating that the 28 signature miRNAs span multiple low-correlation regulatory modules rather than forming a single co-regulated cluster, supporting their joint predictive value.

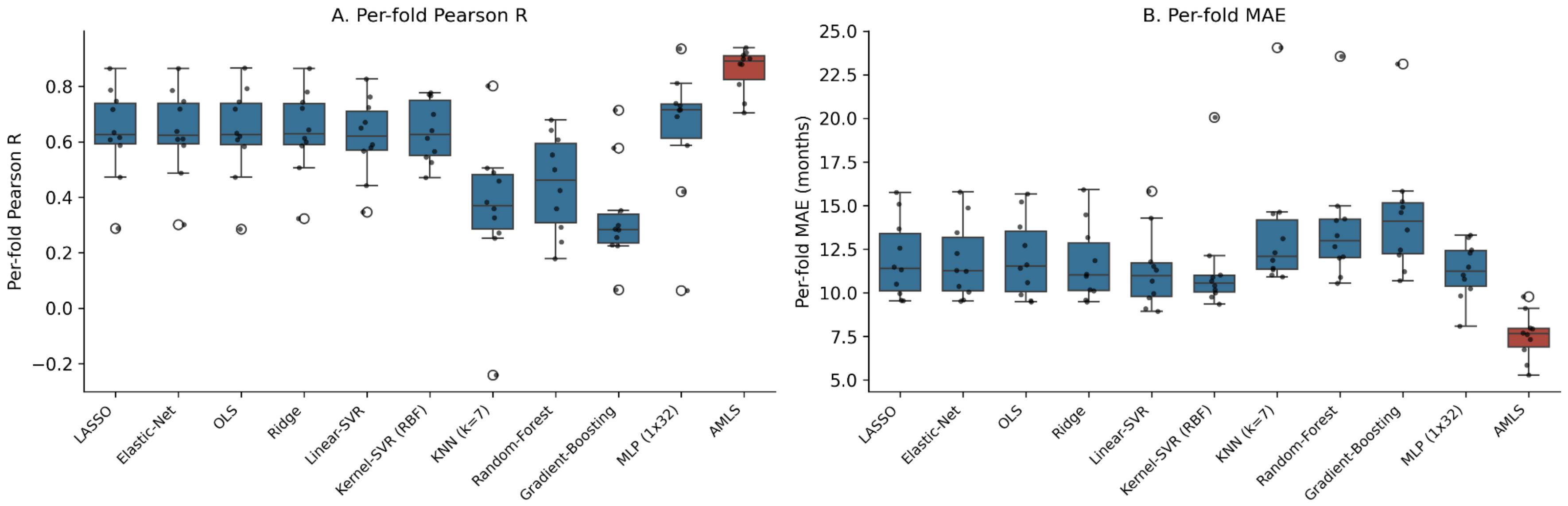


**Figure S5. Per-fold dispersion of model performance.** Box-and-strip plots of per-fold Pearson R (A) and MAE (B) across the 10 outer cross-validation folds for each model on its best feature scenario. AMLS occupies the upper envelope of R and the lower envelope of MAE in every fold.

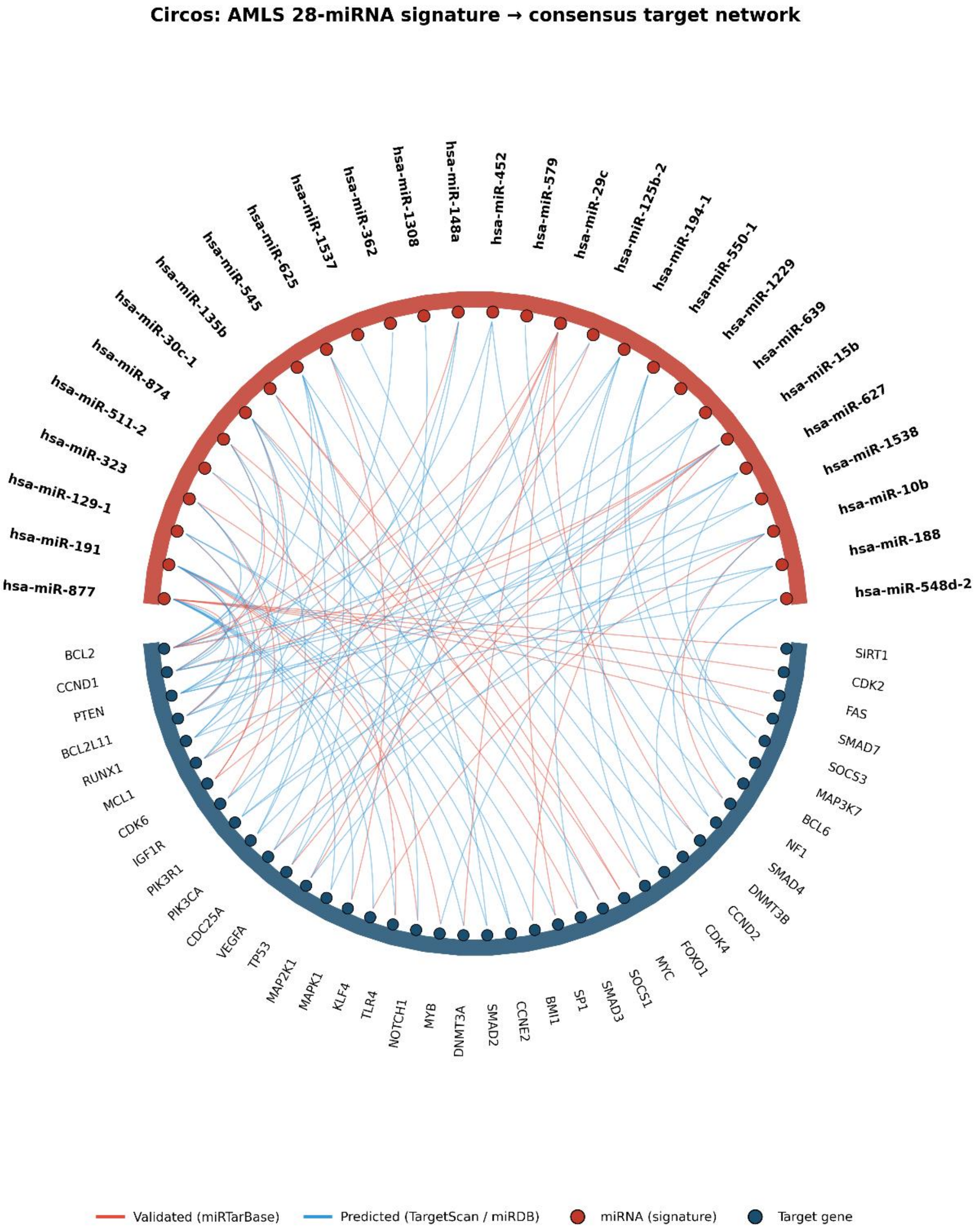


**Figure S6. Detailed miRNA-target circos plot.** Single-ring circos with every miRNA-target chord colored by evidence (validated = red, predicted = blue). High-resolution complement to main Figure 5.

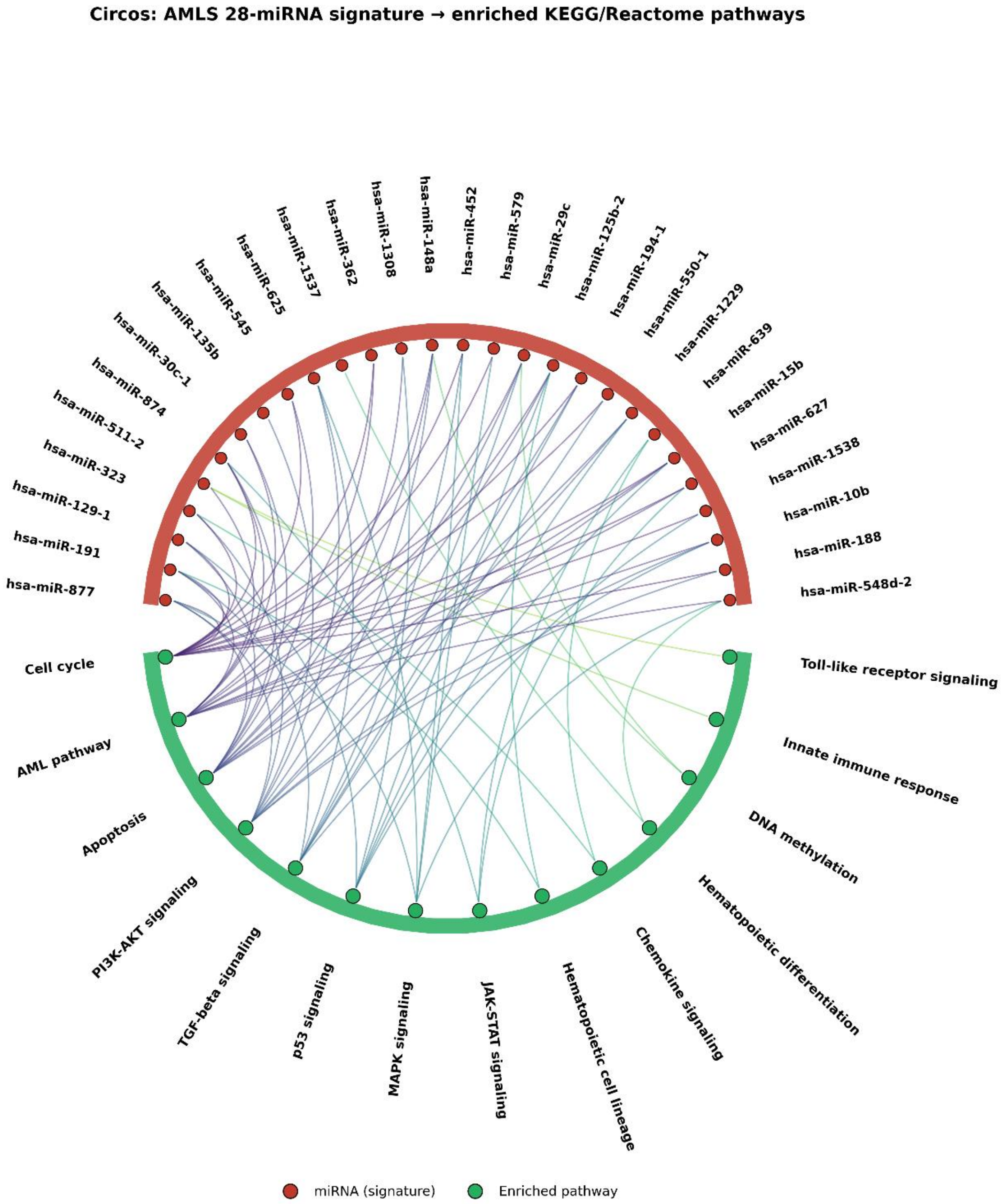


**Figure S7. miRNA-pathway circos plot.** Direct miRNA-to-pathway chord plot — alternative view that emphasizes pathway membership without showing intermediate target genes. Each pathway is color-coded with the viridis colour scale. miR-29c, miR-148a, miR-125b and miR-15b are the most central nodes.

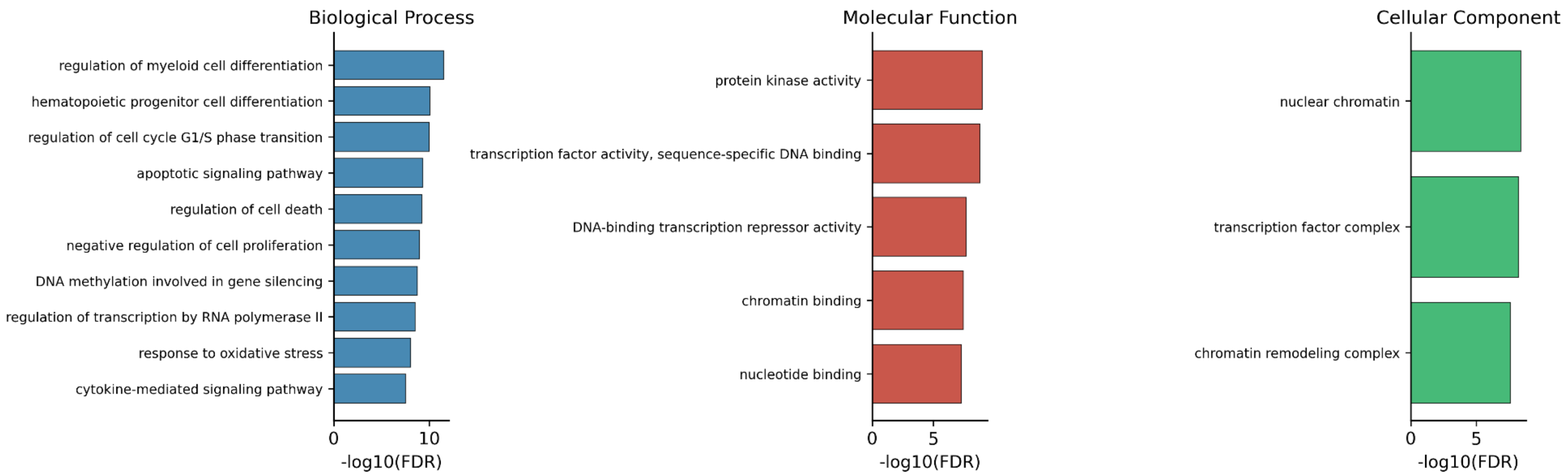


**Figure S8. Gene Ontology enrichment of consensus miRNA targets.** Biological Process (left), Molecular Function (center) and Cellular Component (right) over-representation analyses by hypergeometric test with Benjamini–Hochberg correction. Significant terms (FDR < 0.05) are shown. Top BPs include myeloid-cell differentiation, G1/S regulation, apoptotic signaling and DNA methylation involved in gene silencing.

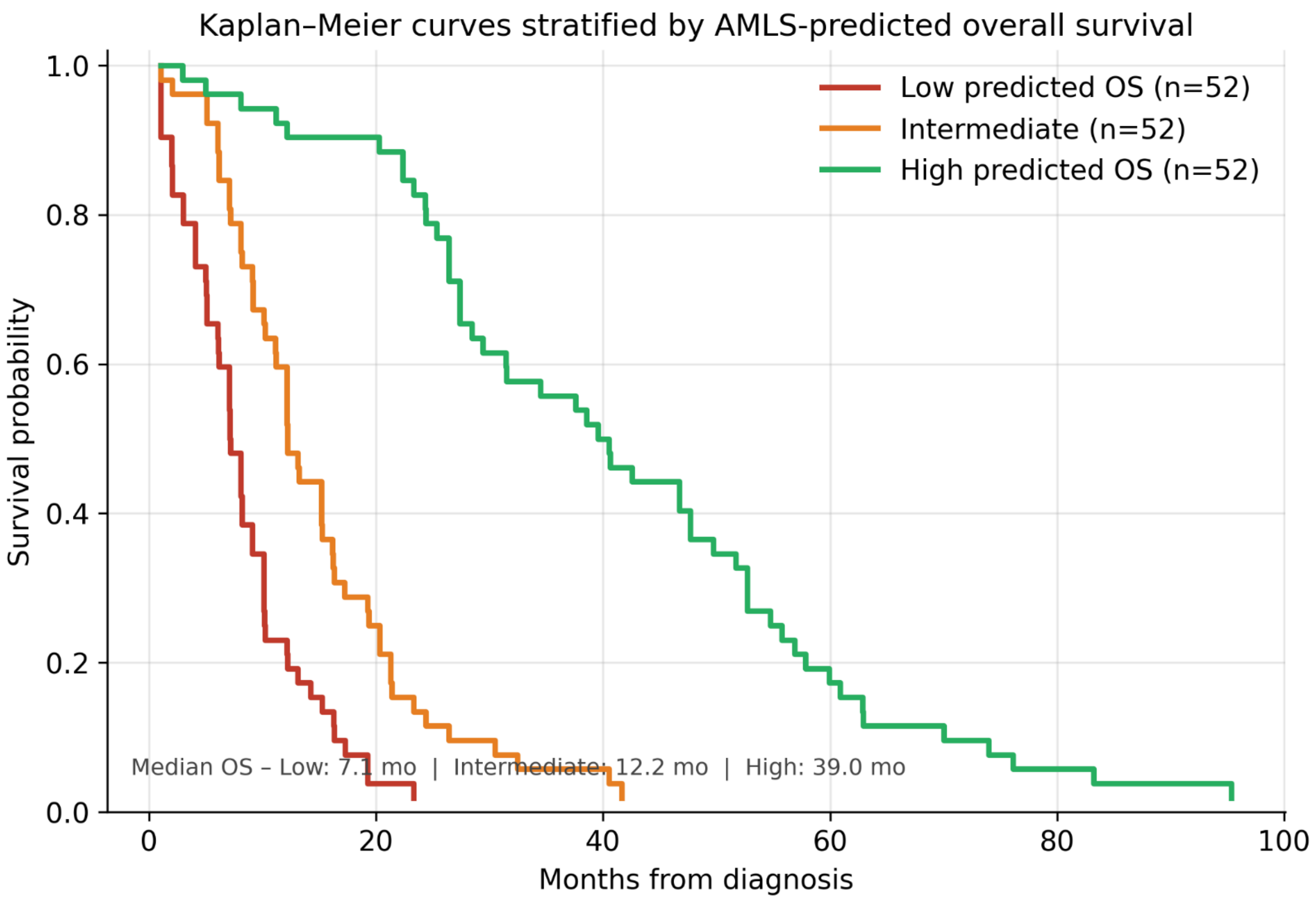


**Figure S9.** Kaplan–Meier curves stratified by tertiles of AMLS-predicted overall survival. Median observed OS: low = 7.1 months, intermediate = 12.2 months, high = 39.0 months.
